# Systematics and geographical distribution of *Galba* species, a group of cryptic and worldwide freshwater snails

**DOI:** 10.1101/647867

**Authors:** Pilar Alda, Manon Lounnas, Antonio A. Vázquez, Rolando Ayaqui, Manuel Calvopiña, Maritza Celi-Erazo, Robert T. Dillon, Luisa Carolina González Ramírez, Eric S. Loker, Jenny Muzzio-Aroca, Alberto Orlando Nárvaez, Oscar Noya, Andrés Esteban Pereira, Luiggi Martini Robles, Richar Rodríguez-Hidalgo, Nelson Uribe, Patrice David, Philippe Jarne, Jean-Pierre Pointier, Sylvie Hurtrez-Boussès

**Affiliations:** Laboratorio de Zoología de Invertebrados I, Departamento de Biología, Bioquímica y, Farmacia, Universidad Nacional del Sur. San Juan No. 670, B8000ICN Bahía Blanca, Buenos Aires, Argentina; MIVEGEC, University of Montpellier, CNRS, IRD, Montpellier, France; Consejo Nacional de Investigaciones Científicas y Técnicas (CONICET), Argentina; Laboratory of Malacology, Institute of Tropical Medicine “Pedro Kourí”, Autopista Novia del Mediodía km 6, La Habana, Cuba; Departamento de Microbiología y Patología, Facultad de Medicina, Universidad Nacional de San Agustín de Arequipa, Peru; Carrera de Medicina, Facultad de Ciencias de la Salud, Universidad de Las Américas, Quito, Ecuador; Instituto de Investigación en Salud Pública y Zoonosis - CIZ, Universidad Central de Ecuador, Quito, Ecuador; Freshwater Gastropods of North America Project, P.O. Box 31532, Charleston, SC 29417, USA; Grupo de Investigación “Análisis de Muestras Biológicas y Forenses” Laboratorio Clínico, Facultad de Ciencias de la Salud, Universidad Nacional de Chimborazo, Ecuador; Center for Evolutionary and Theoretical Immunology, Department of Biology, University of New Mexico, Albuquerque, NM87131, USA; Instituto Nacional de Investigación en Salud Pública INSPI, Guayaquil, Ecuador; Universidad Agraria del Ecuador, Facultad de Medicina Veterinaria y Zootecnia, Guayaquil, Ecuador; Sección de Biohelmintiasis, Instituto de Medicina Tropical, Facultad de Medicina, Universidad Central de Venezuela. Centro para Estudios Sobre Malaria, Instituto de Altos Estudios “Dr. Arnoldo Gabaldón”-Instituto Nacional de Higiene “Rafael Rangel” del Ministerio del Poder Popular para la Salud. Caracas, Venezuela; Grupo de Investigación en Epidemiología Molecular (GIEM), Escuela de Microbiología, Facultad de Salud, Universidad Industrial de Santander, Bucaramanga, Colombia; Laboratorio de Parasitologia Luiggi Martini y colaboradores, Guayaquil, Ecuador; Facultad de Medicina Veterinaria y Zootecnia, Universidad Central del Ecuador, Quito, Ecuador; Centre d’Ecologie Fonctionnelle et d’Evolution, UMR 5175, CNRS – Université de Montpellier – Université Paul Valéry Montpellier – EPHE - IRD, 1919 route de Mende, 34293, Montpellier Cedex 5, France; PSL Research University, USR 3278 CNRS–EPHE, CRIOBE Université de Perpignan, Perpignan, France; Département de Biologie–Ecologie, Faculté des Sciences, Université Montpellier Montpellier, France

**Keywords:** systematics, distribution, America, Lymnaeidae, *Galba*, vector snails, biological invasions

## Abstract

Cryptic species can present a significant challenge to the application of systematic and biogeographic principles, especially if they are invasive or transmit parasites or pathogens. Detecting cryptic species requires a pluralistic approach in which molecular markers facilitate the detection of coherent taxonomic units that can then be analyzed using various traits (e.g., internal morphology) and crosses. In asexual or self-fertilizing species, the latter criteria are of limited use. We studied a group of cryptic freshwater snails (genus *Galba)* from the family Lymnaeidae that have invaded almost all continents, reproducing mainly by self-fertilization and transmitting liver flukes to humans and livestock. We aim to clarify the systematics, distribution and phylogenetic relationships of these species with an integrative approach that includes morphology (shell and reproductive anatomy), molecular markers, wide-scale sampling across America, and data retrieved from GenBank (to include Old World samples). Our phylogenetic analysis suggests that the genus *Galba* originated ca. 22 Myr ago and today comprises six clusters of species. Four of these clusters (*G*. *truncatula*, *G*. *cubensis*/*viator*, *G*. *humilis* and *G*. *schirazensis*) are morphologically cryptic and constitute species or species complexes with wide variation in their genetic diversity, geographic distribution and invasiveness. The other two clusters constitute a single species or a species complex (*Galba cousini*/*meridensis*) that demonstrate more geographically restricted distributions and exhibit an alternative morphology more phylogenetically derived than the cryptic one. Further genetic studies are required to clarify the status of both *G. cousini*/*meridensis* and *G*. *cubensis/viator*. We emphasize that no *Galba* species should be identified without molecular markers and that additional sampling is required, especially in North America, Eurasia and Africa to clarify remaining questions in systematics and biogeography. We also discuss several hypotheses that can explain crypsis in *Galba*, such as convergence and morphological stasis, and hypothesize a role for stabilizing selection in amphibious (rather than strictly freshwater) habitats.

## Introduction

Cryptic species are groups of populations in which the intrapopulation phenotypic variance exceeds interspecific phenotypic variance (see review in Bickford et al., 2007; Fišer et al., 2018; Struck et al., 2018). They have been described in all taxonomic kingdoms, although cryptic species seem to be more often reported in animals (Struck et al. 2018). Their frequency might reach a bit less than 1% of all animal species, an estimate that should be considered quite approximate given our lack of knowledge on the species totals. They also seem to be homogeneously distributed among taxa and biogeographical regions (Pfenninger and Schwenk 2007), although it has been suggested that caves and subterranean habitats may promote cryptic speciation. Such isolated, dark, low-energy habitats promote diversification but constrain morphological differentiation since very specialized adaptations are needed to survive in them (Katz et al. 2018). Four hypotheses have been proposed for the origin of cryptic species (Fig. S1; Bickford et al., 2007; Fišer et al., 2018; Struck et al., 2018): (1) recent divergence, when distinguishing traits have not as yet accumulated (e.g. cave fish, Niemiller et al. 2012); (2) parallelism, where independent phenotypic traits evolve in different taxa from a similar and shared ancestral trait (Struck et al. 2018); (3) convergence, where more distantly related species evolve from dissimilar ancestors (e.g. sea stars, Zulliger and Lessios, 2010), and (4) morphological stasis, where species remain similar over long periods of time constrained by limited genetic variation or stabilizing selection (e.g., Gomez et al., 2004; Struck et al., 2018).

Although interesting as models for study of the speciation process (Coyne and Orr 2004; De Queiroz 2007), cryptic species are problematic from two human perspectives: biological invasion and disease transmission. Cryptic species may exhibit wide differences in invasive ability and impact on invaded ecosystems and communities (Fang et al. 2014). An accurate identification at the species level is required in such situations to track invasions and mitigate any harmful consequences (Kolar and Lodge 2001; Dunn and Hatcher 2015; Jarić et al. 2019). Cryptic species may also exhibit ć differences in disease transmission. This is the case in the *Anopheles gambiae* complex which includes the most important vectors of malaria in Africa (Stevenson and Norris 2016). Some members of the complex are broadly zoophilic, while others feed more strictly on humans. Accurate species identification is required for effective mosquito control.

Snails, especially freshwater ones, are an interesting group for addressing biogeographic issues in cryptic species. Although taxonomists have increasingly used molecular markers over the last decades to identify snail species (Dayrat et al. 2011), morphological characters, especially shell shape and sculpture, remain widely relied upon—often resulting in large numbers of synonymous species (Qian et al. 2012). The shells of some mollusk populations show significant phenotypic plasticity, however, in reaction to temperature, pollution or predation (Bourdeau et al. 2015). Others stay stable for millions of years (e.g., Weigand et al., 2011; Weiss et al., 2018). This has resulted both in the proliferation of species names and descriptions (e.g., Taylor, 2003), most of which are invalid (Jarne et al. 2010; Dillon et al. 2011), as well as the misidentification of valid species, yielding errors in the assessment of species invasion ability and distributional range (e.g., Pfenninger et al., 2006; Rama Rao et al., 2018). For example, an invasive Asian clam of the genus *Sinanodonta* has been overlooked in Russia because it is morphologically indistinguishable from another invasive Asian clam (Bespalaya et al. 2018). Such problems call for an integrated approach to gastropod systematics and biogeography in which phenotypic traits are studied together with appropriate molecular tools (Dayrat 2005).

Here we focus on small-bodied basommatophoran pulmonate snails of the genus *Galba* (Hygrophila, Lymnaeidae), common inhabitants of unstable freshwater habitats worldwide. Baker (1911) recognized 30 species and subspecies in the subgenera *Galba* (*s*.*s*.) and *Simpsonia* in North America, on the basis of minor shell morphological variation, where Hubendick (1951) suggested that as few as four biological species might be valid: *humilis*, *truncatula*, *cubensis* and *bulimoides*. The more recent work of Burch (1982) proposed 22 North American species in the genus *Fossaria*, which we here consider a junior synonym of *Galba*. In South America, Hubendick (1951) recognized only two species of small-bodied, amphibious lymnaeids (*viator* and *cousini*), but more recent work based on molecular approaches distinguishes seven species (*viator*, *schirazensis*, *cousini*, *neotropica*, *meridensis*, *truncatula* and *cubensis*; (Bargues et al. 2007, 2011b, 2011a; Correa et al. 2010, 2011; Lounnas et al. 2017, 2018).

Despite an estimated divergence time on the order of 20 Myr based on genomic data (Burgarella et al. 2015), most of the nominal species of *Galba* share a similar shell morphology and internal anatomy, as well as common levels of phenotypic plasticity in shell, anatomy, and life history (Samadi et al. 2000; Correa et al. 2011). The exceptions are *Galba cousini* (Paraense 1995) and *Galba meridensis* (Bargues et al. 2011b), which are mutually similar but morphologically distinctive. These two groups are cryptic species; they are often misidentified and often confused (Correa et al. 2010; Bargues et al. 2011a).

A further challenge is that crossing cannot be used to distinguish species (Coyne and Orr 2004), as has been done in other freshwater snails (e.g., *Physa* species, Dillon et al., 2011), since *Galba* populations primarily reproduce by self-fertilization (Meunier et al. 2004; Bargues et al. 2011a; Lounnas et al. 2017, 2018). Moreover, at least two species, *G*. *schirazensis* and *G*. *truncatula*, have been shown to be extremely efficient anthropogenic invaders, muddling our knowledge of species distribution. Populations of *G. truncatula*, probably from Eurasia, have invaded South America, especially the Bolivian Altiplano (Meunier et al. 2004). This is of special concern since *Galba* populations are the main vectors of the liver fluke *Fasciola hepatica* which causes fasciolosis in both livestock and humans (Mas-Coma et al. 2005) and transmission efficiency and invasion ability differ among species (Vázquez et al. 2018).

Worldwide, the geographic distribution of *Galba* species is poorly known. The dubious character of records based on morphological identification leaves us with a small sample of molecular studies (Correa et al. 2010, 2011; Bargues et al. 2011b, 2011a, 2012; Lounnas et al. 2017, 2018) upon which to base a very large-scale phenomenon. So, our objectives here are to characterize the geographic distribution of *Galba* species at continental scale, based on an extensive sampling over America, to reconstruct the genus phylogeny to delimit species, and to explore the origin of crypsis. We aim to delineate species—the real scientific challenge of integrative taxonomy (Dayrat 2005)—noting that in practice *Galba* species are very difficult to delineate due to its crypsis, wide geographical distribution, and mating system. Previous studies reconstructing *Galba* phylogeny have used single genes and analyzed fewer than 10 sequences per species, failing to account for the wide geographic distribution of this genus (Correa et al. 2010, 2011; Bargues et al. 2011c, 2011a; Standley et al. 2013). Here we employ morphological and molecular markers (microsatellite loci and DNA sequences from two genes) to study more than 1,700 individual *Galba* from 161 sites. Our data set was augmented with a complete sample of all the *Galba* DNA sequences available in GenBank and multiple phylogenetic analyses conducted. We used both gene trees and multispecies coalescent models to shed light on the phylogenetic relationships and on the origin of crypsis in the genus *Galba*.

## Materials and methods

### Snail sampling and species identification

We conducted simple searches for *Galba* populations in suitable habitats throughout the New World over a span of two decades, 1998–2017. Overall, *Galba* populations were detected in 161 sites and 1,722 individuals were sampled from nine countries: Argentina, Canada, Colombia, Cuba, Ecuador, France (French Guiana, Guadeloupe and Martinique), Peru, Venezuela, and USA (Table S1). *Galba* populations have previously been reported from some of these sites in Venezuela and Ecuador by the authors (Pointier 2015; Orlando Narváez et al. 2017).

**Table 1.** Type localities of *Galba* species and sequences of individuals recovered in those localities. References mentioned are the ones in which sequences were provided and not in which species were described, except for articles describing *Galba meridensis* and *Galba neotropica* were the authors also provided sequences (Bargues et al. 2007, 2011b). The type localities for *Galba cubensis* and *Galba viator* were here restricted because original description of the species did not provide a locality, authors simply stated “Cuba” and “Patagonia”, respectively (D’Orbigny 1835; Pfeiffer 1839).

| Species | Country | Locality | Coordinates | ITS1 | ITS2 | 16S | COI | Reference |
| --- | --- | --- | --- | --- | --- | --- | --- | --- |
| *Galba cousini* | Ecuador | Chanchu-Yacu | 00°18'55"S<br>78°34'02"W | FN598157 | FN598153 | - | FN598161 | Bargues et al. 2011b |
| *Galba cubensis* | Cuba | Vaqueria 21 | 23°01'N<br>82°32'W | AM412226 | AM412223 | - | AM494009 | Bargues et al. 2007 |
| *Galba humilis* | USA | Owego, New York | 42°06'01"N<br>76°15'40"W | FN182193,<br>FN182194 | FN182191,<br>FN182192 | - | FN182197,<br>FN182198 | Correa et al. 2011 |
| *Galba meridensis* | Venezuela | Laguna Mucubaji (Mérida) | 08°47'52"N<br>70°49'32"W | FN598159 | FN598154 | - | FN598164 | Bargues et al. 2011b |
| *Galba neotropica* | Peru | Lima, Rio Rimac | 12°02'S<br>76°56'–<br>77°08'W | AM412228 | AM412225 | - | AM494008 | Bargues et al. 2007 |
| *Galba schirazensis* | Iran | Taleb Abad river, Bandar Anzali, Gilan province | 37°27'46"N<br>49°37'07"E | JF272603 | JF272601 | JF272605 | JF272607 | Bargues et al. 2011a |
| *Galba truncatula* | Germany | Thuringia, Erfurt-Bindersleben | ND | - | - | - | EU818799 | Albrecht et al. 2008 |
| *Galba viator* | Argentina | Frias | 40°14' S<br>64°10' W | JN614428 | HQ283265,<br>JN614465 | - | JN614397,<br>JN614398 | Correa et al. 2011 |

In most cases, we discovered our *Galba* populations in unstable habitats subject to frequent flooding and droughts. The sampled habitats were characterized as brook, irrigation canal, ditch, oxbow lake, pond, marsh, lake, tank, rice field, and river. Individual snails were often collected above the water line, consistent with their amphibious habit. Some sites were visited up to five times. Geographic coordinates were recorded for most sites.

After collection, individuals were placed in water at 70 °C for 30–45 s. This procedure allows fixation of individuals without contraction of soft parts and facilitates a proper study of snail internal anatomy. The body was carefully withdrawn from the shell using forceps and both body and shell stored in 70% ethanol until morphological and DNA analyses (Pointier et al. 2004).

Species were characterized using a three-step procedure involving both morphological and molecular markers (Fig. 1). Step 1 was an analysis of shell morphology and reproductive anatomy. In step 2, we used a molecular tool that enables us to distinguish *G*. *cubensis*, *G*. *schirazensis*, and *G*. *truncatula* (Alda et al. 2018). In step 3, we sequenced mitochondrial and nuclear genes in individuals for which no PCR amplification product was obtained in step 2. DNA from some of those individuals identified in steps 1 and 2 were also sequenced in order to reconstruct the phylogeny of *Galba*. Note that this three-step approach is less expensive than an approach based on simply sequencing the same genes in all individuals.

**Figure 1.**
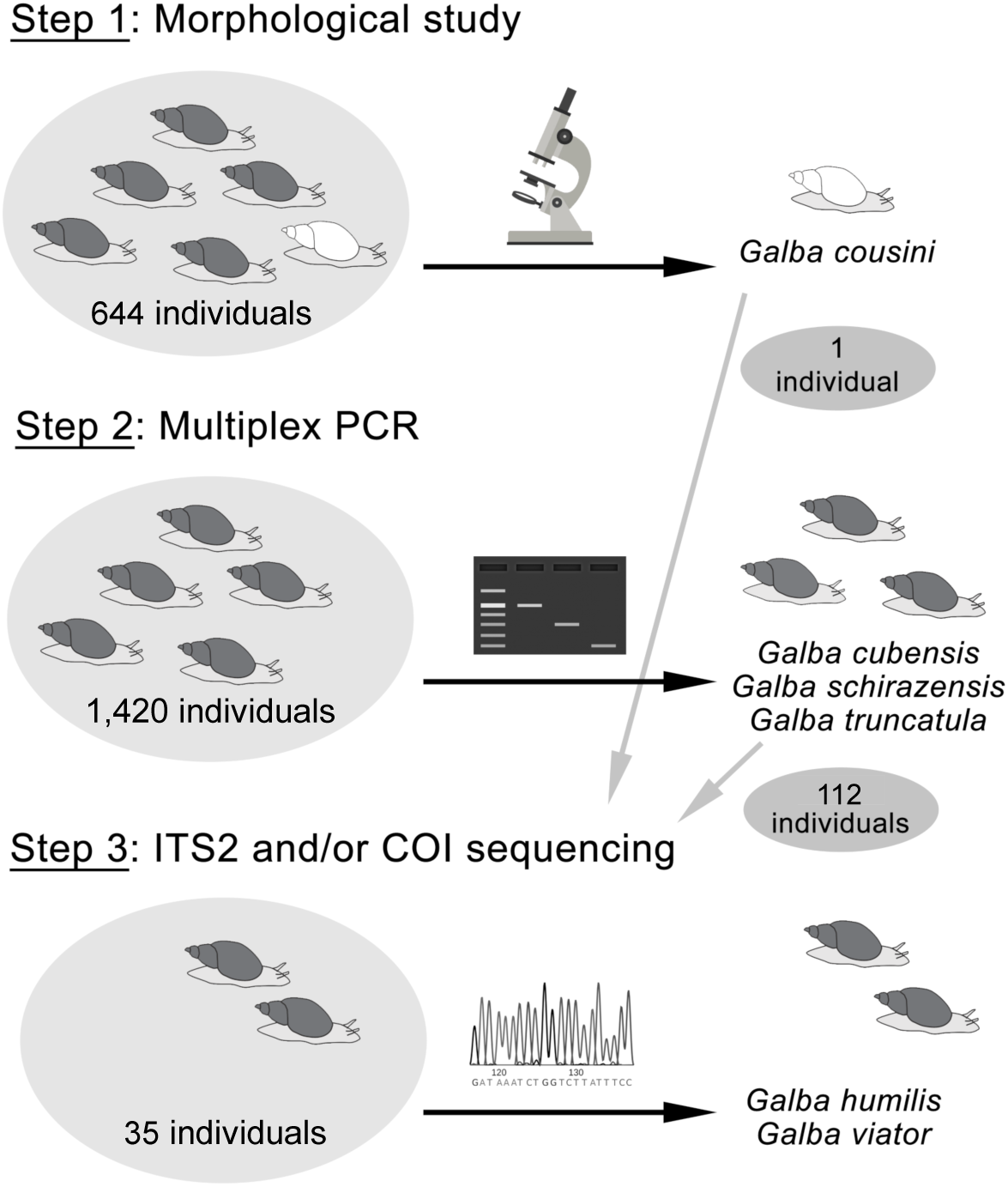
The three-step procedure followed to identify the 1,722 individuals of *Galba* species. The number of individuals identified at each step is indicated in the left and the species identified are indicated on the right. In Step 1, we photographed the shell and dissected three to five adult snails from each of the 166 sites. Fragments of the ITS2 and COI genes were sequenced in 146 individuals: *Galba cousini*/*meridensis* (1), *Galba cubensis* (41), *Galba schirazensis* (41), *Galba truncatula* (30), *Galba humilis* (34) and *Galba viator* (1).

#### Step 1: morphology of the shell and of internal organs

We photographed the shell of three to five adult snails from each site and dissected their body under a stereoscopic microscope. We drew the anatomy of the penial complex, prostate, and renal tube using a camera lucida attachment (Pointier et al. 2004). We did not record any morphological measurements or perform any quantitative tests because Correa et al. (2011) have shown that cryptic *Galba* species cannot be delimited by such means. Our observations were qualitative only (Fig. S2).

**Figure 2.**
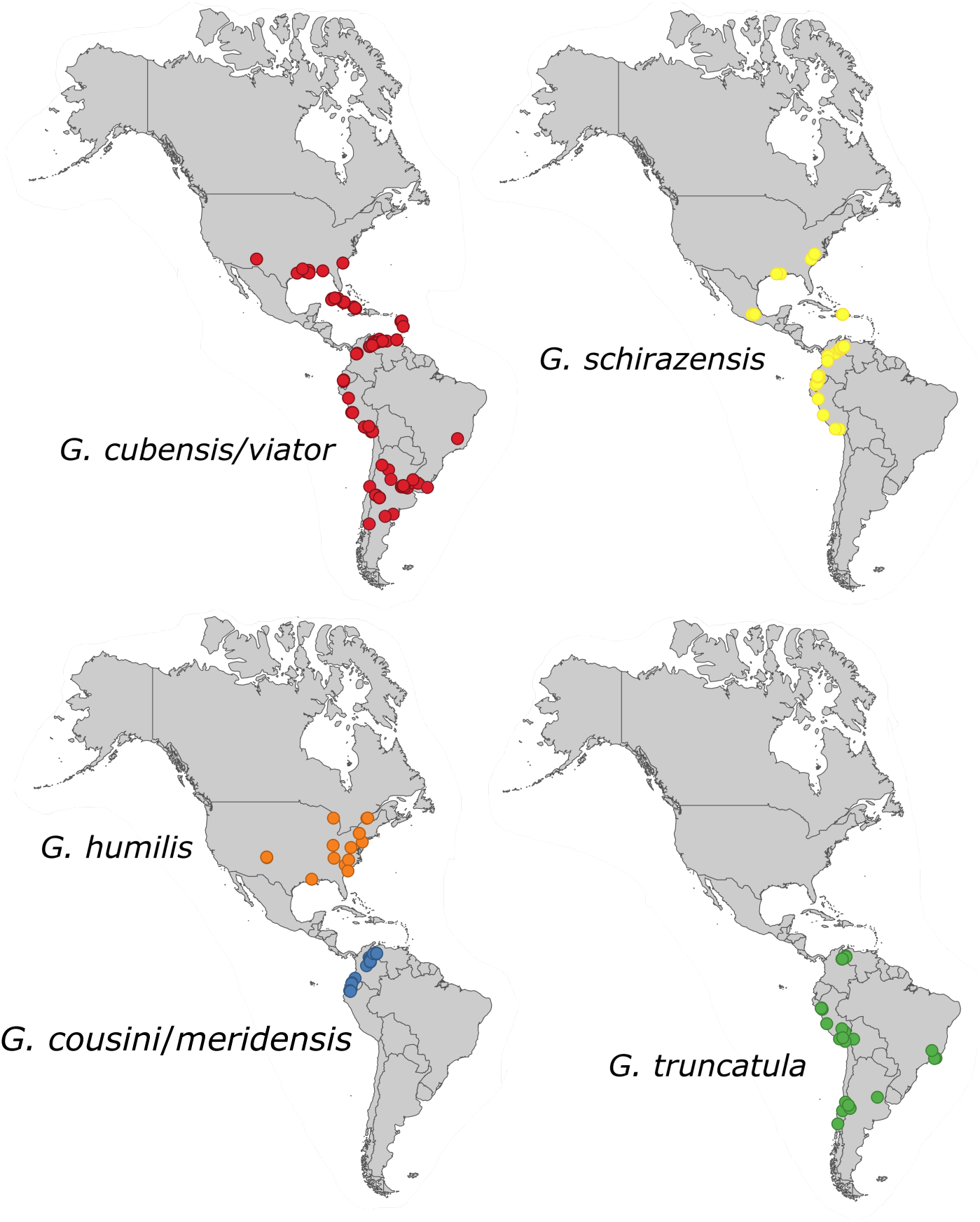
Geographic distribution of *Galba* species in America, based on molecular identification. Coordinates are given in Tables S1–S2. The sites Perdriel (Argentina), Batallas and Tambillo (Bolivia), Ontario (Canada), San Rafael (Mexico), Canal Salinas (Puerto Rico) are not represented since coordinates are missing in the original publications.

#### Step 2: multiplex PCR of microsatellite loci

We applied the multiplex PCR test designed by Alda et al. (2018) to all the 1,420 individuals that were not distinguishable based on shell and reproductive anatomy. This method is based on species-specific primers amplifying three microsatellite loci (one each per targeted species) and producing band sizes that are specific to these species (179–200 pb in *G*. *cubensis*, 227–232 pb in *G*. *schirazensis* and 111–129 pb in *G*. *truncatula*). DNA was extracted using a Chelex protocol following Estoup and Martin (1996) as adapted for 96-well plates. Methods for DNA amplification and electrophoretic resolution followed Alda et al. (2018).

#### Step 3: identification by sequencing

We amplified the internal transcribed spacer 2 (ITS2) and the cytochrome oxidase subunit 1 (COI) genes in 35 individuals sampled from 15 sites in Argentina, Canada and the USA (1 to 5 per population) where at least some individuals did not demonstrate an amplification product in step 2, using the method of Lounnas et al. (2017, 2018). We also amplified ITS2 and COI in 112 individuals that did return an amplification product in step 2, including one individual identified as *G*. *cousini*/*meridensis*. To supplement these results, we also amplified the internal transcribed spacer 1 (ITS1) and the ARN ribosomal 16S in two individuals of *G. cubensis* from Bosque del Apache (USA) and in one individual of *G. cousini*/*meridensis* from Ecuador (Table S1). With this approach, we obtained at least one sequence from each hypothetical species represented by the four genes and used them to delimit species. The total number of individuals amplified was 151.

We used the primers NEWS2 (forward) 5’ TGTGTCGATGAAGAACGCAG 3’ and ITS2-RIXO (reverse) 5’ TTCTATGCTTAAATTCAGGGG 3’ to amplify ITS2; Lim1657 (forward) 5’ CTGCCCTTTGTACACACCG 3’ and ITS1-RIXO 5’ TGGCTGCGTTCTTCATCG 3’ to amplify ITS1 (Almeyda-Artigas et al. 2000); LCOI490 (forward) 5’ GGTCAACAAATCATAAAGATATTGG 3’ and HCO2198 (reverse) 5’ TAAACTTCAGGGTGACCAAAAAATCA 3’ to amplify COI (Folmer et al. 1994) and forward 5’ CGCCTGTTTATCAAAAACAT 3’ and reverse 5’ CCGGTCTGAACTCAGATCACGT 3’ to amplify 16S (Remigio and Blair 1997). In all cases, PCR amplification was performed in a total volume of 25 µl containing 12.5 µl of Taq PCR Master Mix (Qiagen), 2.5 µl of each primer (10 mM) and 2 µl of DNA in an Eppendorf Thermal Cycler with an initial denaturation step at 95 °C for 15 minutes; followed by 35 cycles of denaturation at 95 °C for 30 seconds, annealing at 50 °C for one minute, extension at 72 °C for one minute; and a final elongation at 60 °C for 30 minutes. The presence and size of amplification products were electrophoretically confirmed in 1% agarose gels stained with EZ-Vision. DNA sequencing was performed by Eurofins Genomics (Ebersberg, Germany) using PCR-amplified products as templates. All sequences were uploaded to GenBank (Table S1) and assigned to a species using the phylogenetic reconstruction.

### Type localities

Because of the longstanding confusion and uncertainty regarding the systematics of the genus *Galba* worldwide, it is especially important to establish standard populations, against which unknown populations can be compared. Type localities were specified by the authors of all the more recently-described species, such as *neotropica* (Bargues et al. 2007) and *meridensis* (Bargues et al. 2011b), and others have been established by subsequent use, for example *schirazensis* (Bargues et al. 2011a). But in his original description of *Limnaeus viator*, D’Orbigny (1835) simply stated “Patagonia.” And Pfeiffer (1839) gave no locality data for his *Limnaeus cubensis* at all, beyond “Cuba”. In such circumstances, the ICZN code provides that subsequent authors may restrict type localities to some more precise spot “within the known range of the taxon”. Type localities for all eight of the widely-recognized species in the genus *Galba*, either as originally stated or as subsequently restricted, are listed in Table 1. COI sequences from samples of all the populations inhabiting these localities have been previously uploaded to GenBank, and most have ITS1, ITS2 or 16S sequences available as well.

### Retrieving data on *Galba* spp. distribution from published work

We searched the literature and GenBank for sequence data at four genes (COI, ITS1, ITS2, and 16S) apparently attributable to lymnaeids of the genus *Galba*. Coordinates were provided for most sites by the authors. When coordinates were not provided, we inferred them from the locality data using GoogleEarth. We found 132 New World sites in which *Galba* species have been molecularly characterized (Table S2), and 45 sites in the Old World (Table S3, Fig. S3). The specific nomina attributed to these sequences by their depositors in GenBank were 157 *truncatula*, 152 *schirazensis*, 70 *neotropica*, 57 *cubensis*, 44 *viator*, 20 *cousini*, 9 *humilis*, 6 *meridensis*, and 2 others. This corresponds to 166 COI, 163 ITS2, 118 ITS1, and 70 16S sequences.

**Figure 3.**
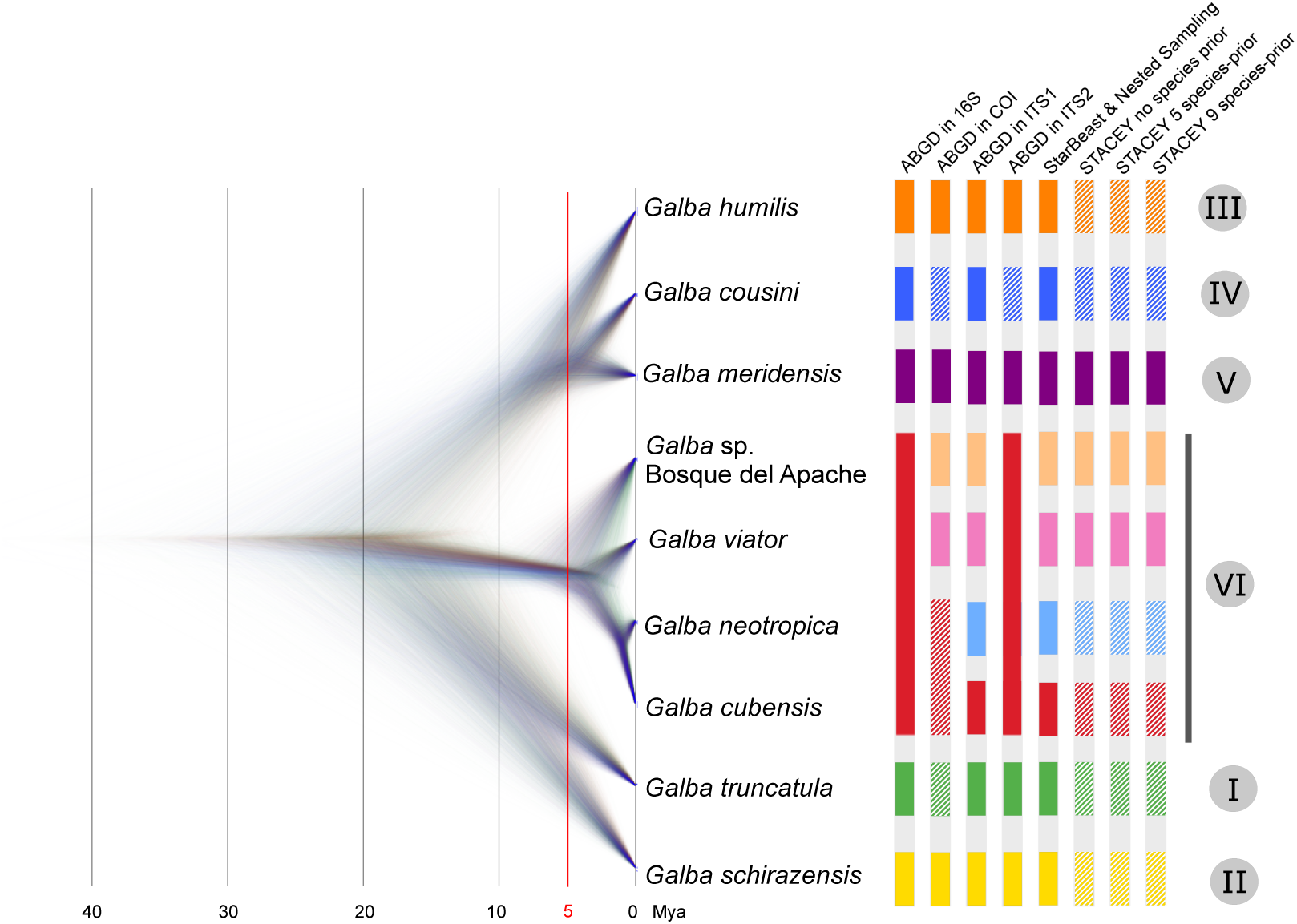
Time-calibrated phylogenetic hypotheses of the model that best approximate species status in *Galba* and species-delimitation methods. The trees showed are the most probable topologies based on the multispecies tree model that showed the highest Bayes Factor in StarBeast2 (visualized in Densitree), as a function of time (in Mya). Greater topological agreement is visualized by a higher density of trees (on the left), whereas uncertainty in the height and distribution of nodes are represented by increased transparency. The most common topologies are shown in blue, the second most common topologies in red and the third in green. The six clusters built with individual gene trees are indicated on the extreme right. The number of species recovered varied with eight approaches (ABGD with each gene, StarBeast with all genes, STACEY with a variable number of species included as prior). For each approach, the colored bars represent different species. Bars were striped when the groups included more than one species. Clusters as appeared in text are shown in roman numbers. Scale bar represents branch length expressed as number of substitutions per site.

### Phylogenetic and ancestral reconstruction study

Phylogenetic analyses were conducted on the ITS2 and COI sequences obtained in this study, together with ITS2, COI, ITS1, and 16S sequences retrieved from GenBank. Most of these sequences have been previously published (Tables S2–S3). Ultimately, we included 796 sequences in our analyses: 90 for 16S, 251 for COI, 122 for ITS1, and 333 for ITS2. Some GenBank accession numbers appear more than once in Tables S2– S3 because individuals with identical sequences have been registered under the same GenBank accession number.

Alignment was performed individually for each gene using MAFFT (Katoh and Standley 2013). Ambiguously aligned sites were excluded using GBLOCKS with default settings for a less stringent selection (Castresana 2000). The number of positions in the final sequences was 412 for 16S (83% of the original 493 positions), 609 for COI (86% of 707), 435 for ITS1 (48% of 888), and 333 for ITS2 (23% of 1,429). We examined levels of saturation for each gene and for the first and second *versus* third codon positions of COI using DAMBE (Xia 2017). We did not find evidence of saturation in the four genes analyzed, including all codon positions of COI. We partitioned codon positions and unlinked substitution models in phylogenetic analyses.

We used Bayesian inference in Beast2 (Bouckaert et al. 2014) for five reasons: (1) to assign sequences to species; (2) to validate, and to mend if necessary, species identity for sequences retrieved from GenBank; (3) to delimit species; (4) to reconstruct the phylogeny and estimate time divergence; and (5) to assess the ancestral phenotypic state of *Galba* species. Because we did not have identical sets of individuals (or populations) across genes it was necessary to build four unlinked gene trees to address questions (1) and (2). The best-fitting models of sequence evolution for each gene were selected using bModelTest (Bouckaert and Drummond 2017). We estimated a model for each COI partition (1^st^, 2^nd^, and 3^rd^ position). The best model describing the evolution of 16S was HKY+G+I, 123424+G+I for COI (1^st^ codon), 121321+G+I for COI (2^nd^ codon), TN93+G+I for COI (3^rd^ codon), 123424+G+I for ITS1, and 121323 for ITS2. We linked trees for the three COI codon partitions. The analyses were run using four gamma categories and a proportion of 0.5 invariant sites. Uncorrelated relaxed-clock models were chosen for all loci. The relative clock mean priors were all lognormal (M = 0, S = 1). We used a birth-death model as priors with lognormal birth and death rates.

These gene trees allowed us to evaluate species names and validate or mend species identity for all sequences, whether obtained in this study or retrieved from GenBank. All the MCMC were run for 200,000,000 generations storing every 20,000 generations. The names of the eight widely-recognized species in the genus *Galba* were ultimately assigned by reference to their type localities. Since gene trees can be erroneously inferred due to incomplete sampling, incomplete lineage sorting or introgression between lineages, we also built haplotype networks for each gene using popART (Leigh and Bryant 2015) and compared them with gene trees.

To address our question (3), species delimitation, we built ten multispecies coalescent tree models in StarBeast2 (Ogilvie et al. 2017) differing in species assignments, some models splitting species to as many as nine while others lumping to as few as five. We assigned the species identity obtained from tasks 1 and 2 to each individual and tested each scenario in turn. We also created an eleventh model in which we separated the populations of *G. viator* from Argentina and from Chile to test whether splitter models showed higher support than lumper models regardless of their biological meaning. Since all the hypothetical species generated in these tree models must be represented by all genes, we removed the sequences from Ethiopia in this analysis.

In all our multispecies coalescent tree models, we used the same site models as for reconstructing the gene trees. We assigned to the mitochondrial loci a gene ploidy of 0.5 and to nuclear loci a gene ploidy of 2.0 (diploid). We used constant population sizes (fixed to 1) and uncorrelated relaxed-clock models for all loci. For nuclear loci, we used the molecular clock rate reported by Coleman and Vacquier (2002) for ITS in bivalves (0.00255 per Myr). For mitochondrial loci, we retained the molecular clock rate estimated by Wilke et al. (2009) for COI in invertebrates (0.0157 per Myr). We used the birth-death model as tree prior with lognormal birth and death rates. We ran each model using Multi-Threaded Nested Sampling analysis with 10 particles, 8 threads, a chain length of 100,000,000 and sub-chain length of 5,000. Then, we compared the trees by computing Bayes factor (BF), a model selection tool that is simple and well-suited for comparing species-delimitation models (Leaché et al. 2014). BF is the difference between the marginal likelihoods of two models: BF = model 1 – model 2. If BF is larger than 1, model 1 is favored, and otherwise model 2 is favored. When BF is between 20 and 150 the support strong and when the BF is above 150 the support is overwhelming (Leaché et al. 2014). We used Nested Sampling implemented in the NS package to calculate the marginal likelihoods necessary to obtain BF and also an estimate of the variance of the marginal likelihoods (Russel et al. 2019).

We also applied two other species-delimitation methods: Automatic Barcode Gap Detection (ABGD; Puillandre et al. 2012) and Species Tree and Classification Estimation in Beast2 (STACEY, Jones 2017). ABGD was run for each gene using the default settings (https://bioinfo.mnhn.fr/abi/public/abgd/). The prior on intraspecific divergence defines the threshold between intra- and interspecific pairwise distances and is iterated from minimum to maximum through ten steps (Puillandre et al. 2012). Given the wide genetic diversity observed within some species (Lounnas et al. 2017), we chose the partition that showed high prior on intraspecific divergence (the penultimate partition).

For the STACEY delimitation method, we used only the ITS2 and COI sequences because these both genes were better represented in the dataset than ITS1 and 16S. Since the hypothetical number of species in STACEY ranges from one to the number of individuals, each of our 113 snail populations was considered as a minimal cluster. The method distinguishes very shallow species divergences with a statistic called “collapseHeight,” which we set to a small value (0.0001) following Jones (2017). Given the results obtained with ABGD and Nested Sampling analyses, we ran three multispecies coalescence models with different collapseWeight parameters: (1) a lumping model with five *Galba* species with 1/X distribution (initial value: 0.975, between 0 and 1), (2) a splitting model with nine species with 1/X distribution (initial value: 0.952 between 0 and 1), and (3) a no-prior-taxonomic model with a Beta distribution (α = 2, β = 2). The initial value of the lumping and splitting models were calculated following Matos-Maravi et al. (2018). All other parameters were set as in the Nested Sampling analysis. We ran the three models for 250,000,000 generations with storing every 25,000 generations.

To address our question (4), regarding species topology and divergence time, we reran the multispecies tree model that showed the highest Bayes Factor in StarBeast2. We used the same parameters as in the Nested Sampling analysis, but we ran the MCMC for a longer time (250,000,000 generations stored every 25,000 generations). The MCMC output was visualized using Tracer (Rambaut et al. 2018) and tree samples summarized by TreeAnnotator (utility program distributed with the Beast package) using a 10% burn-in. The species tree was visualized and edited in FigTree, GIMP (https://www.gimp.org), and DensiTree (Bouckaert and Heled 2014). Some analyses were run in CIPRES Science Gateway (Miller et al. 2012). Note that we did not add an outgroup to root trees because *Galba* monophyly has already been ascertained (Correa et al. 2010, 2011).

Finally, to address our question (5), the ancestral phenotypic state, we applied Bayesian Binary MCMC (BBM, Ronquist and Huelsenbeck 2003), statistical dispersal-vicariance analysis (S-DIVA; Yu et al. 2010), and Statistical dispersal-extinction-cladogenesis (S-DEC; Ree and Smith 2008) in the software Reconstruct Ancestral State in Phylogenies (RASP, Yu et al. 2015). We used the splitting model (scenario A) under default settings. We added two phenotypic states: one for all cryptic species and another one for *G*. *cousini*/*meridensis*.

## Results

### Morphology

Most individuals (N = 1,420 from 133 sites) were not distinguishable based on shell and reproductive anatomy (Fig. S4). The exception was a single group comprising all individuals from *G. cousini* and *G. meridensis* (N = 302). These tended to demonstrate more globose shells with shorter spires, adult sizes in excess of 10 mm standard shell length. *Galba cousini* and *G. meridensis* also differed from the other species in their internal anatomy—a ureter with two distinct flexures, a wider and more ovate prostate, a larger penial complex, and a penis sheath approximately the same length as the preputium (Fig. S4). We did not find any anatomical differences between *G. cousini* and *G. meridensis*, however, comparing individuals from Ecuador, Colombia, and Venezuela (Fig. 2).

### Multiplex PCR of microsatellite loci

DNA from the 1,420 American individuals with similar phenotypes was amplified using the multiplex PCR procedure (step 2 from Fig. 1). We identified 541 individuals of *G*. *cubensis*, 330 of *G*. *schirazensis*, and 349 of *G*. *truncatula* (Table S1; Fig. 2). No amplification was observed in 200 individuals sampled in one site from Argentina and 14 sites from Canada and USA (Table S1).

### Identification by sequencing

Phylogenetic analysis of COI sequences returned six clusters (Fig. S5). Clusters I–V each contained a single type population: *truncatula* (I), *schirazensis* (II), *humilis* (III), *cousini* (IV), and *meridensis* (V). Cluster VI contained the type populations of *cubensis*, *neotropica*, and *viator*. The posterior probabilities (PP) of all clusters were 1.0, except for cluster III (*humilis*, PP = 0.95).

Analysis of 16S, ITS1, and ITS2 sequences confirmed the COI results in almost all respects, although sequences were missing for at least one or two type populations in each tree (Figs. S6–S8). However, we detected discrepancies between the mitochondrial and nuclear gene trees. A striking example are sequences from Bosque del Apache (USA; cluster VI) that exhibited very long branches in the mitochondrial gene trees (Figs. S5–S6) but clustered tightly with the *cubensis* type population and similar populations in the nuclear gene trees (Figs. S7–S8). The mitochondrial sequences from Ethiopia (one per gene) also formed a long branch, although nuclear sequences were not reported in GenBank for that population. Within cluster VI, we found that the *cubensis* and *neotropica* type populations were located in separate subclades for the nuclear genes but clustered together in the mitochondrial trees (Figs. S5–S8).

Gene trees (Figs. S5–S8) and haplotype networks (Figs. S9–S12) showed that genetic diversity varied among the six clusters and four genes. Cluster II (*schirazensis*) showed reduced variation, while cluster VI (*cubensis*, *neotropica*, and *viator*) was larger and more diverse. Mitochondrial genes seemed more diverse than nuclear genes. This observation may be biased, however, by a structural correlation in our data between genes sequenced and regions sampled.

Most of the sequences uploaded to GenBank identified as one of the eight species of *Galba* were accurately clustered into the six clades containing their type populations. However, there were some exceptions. Eight sequences of COI uploaded as *G*. *truncatula* from France appeared in cluster II with the *G*. *schirazensis* type population (Table S3; Fig. S5). We reidentified these sequences as belonging to *G*. *schirazensis*. The COI sequence from Ethiopia, uploaded as *G*. *truncatula*, clustered at the base of *truncatula* clade I with low posterior probability (Fig. S5). The other Ethiopian sequence (16S), also identified as *G*. *truncatula*, did not cluster with any of the *Galba* clades (Fig. S6).

### Species delimitation

Figure 3 illustrates the results obtained using the three species-delimitation methods. The Multi-Threaded Nested Sampling analysis (Fig. S13) suggested that scenario A (nine species) is the best fit to the available data, demonstrating the largest maximum likelihood estimate (Fig. 3; see Table S4 for Likelihoods). BF analysis preferred scenario A over scenario D (current taxonomy) or scenario K, separating populations of *G. viator* from Argentina and from Chile. This suggests that further splitting the phylogeny of *Galba* (here to consider that *G. meridensis* and *G. cousini* are separate species) is not required.

ABGD results varied depending on the gene analyzed (Fig. 3). Nine species (scenario A) were suggested by ITS1, while six species only were returned by our analysis of ITS2 and 16S, with *G. cubensis*, *G. viator*, *G. neotropica*, and *Galba* sp. “Bosque del Apache” lumped. ABGD analysis of the COI gene indicated that *G. viator* and *Galba* sp. “Bosque del Apache” are separate species, but that *G. cubensis* and *G. neotropica* should be lumped together. The ITS2 and COI analyses also suggested that some species (*G. cousini* and *G. truncatula*) might be represented by more than one taxon.

The species-delimitation analysis implemented in STACEY suggested that six of the nine clusters of scenario A might include more than one taxon. The exceptions were *G. viator*, *G. meridensis*, and *Galba* sp. “Bosque del Apache,” the last two species including only one population. Our STACEY results converged towards similar MCMCs regardless of which prior was used for the collapseWeight parameter (Fig. 3).

### Phylogeny, time of divergence, and state reconstruction

Clusters III (*humilis*), IV (*cousini*), and V (*meridensis*) were grouped together in all gene trees returned by our analysis. But inconsistent results among genes were obtained for the remainder of the other clusters identified (Figs. S5–S8). The multilocus multispecies tree returned three major groups: cluster I (*truncatula*) together with II (*schirazensis*); cluster III (*humilis*) together with IV (*cousini*) and V (*meridensis*); and the cluster VI group (*viator*/*cubensis*/*neotropica*/*sp.* “Bosque del Apache”). All groups showed posterior probabilities of 1.0 except for the first one (0.44; Fig. 4). The multilocus multispecies tree visualized in DensiTree (Fig. 3) confirmed that most tree topologies united the clusters into the three major groups outlined above, although some topologies placed clusters differently reflecting the incongruence found among the gene trees.

**Figure 4.**
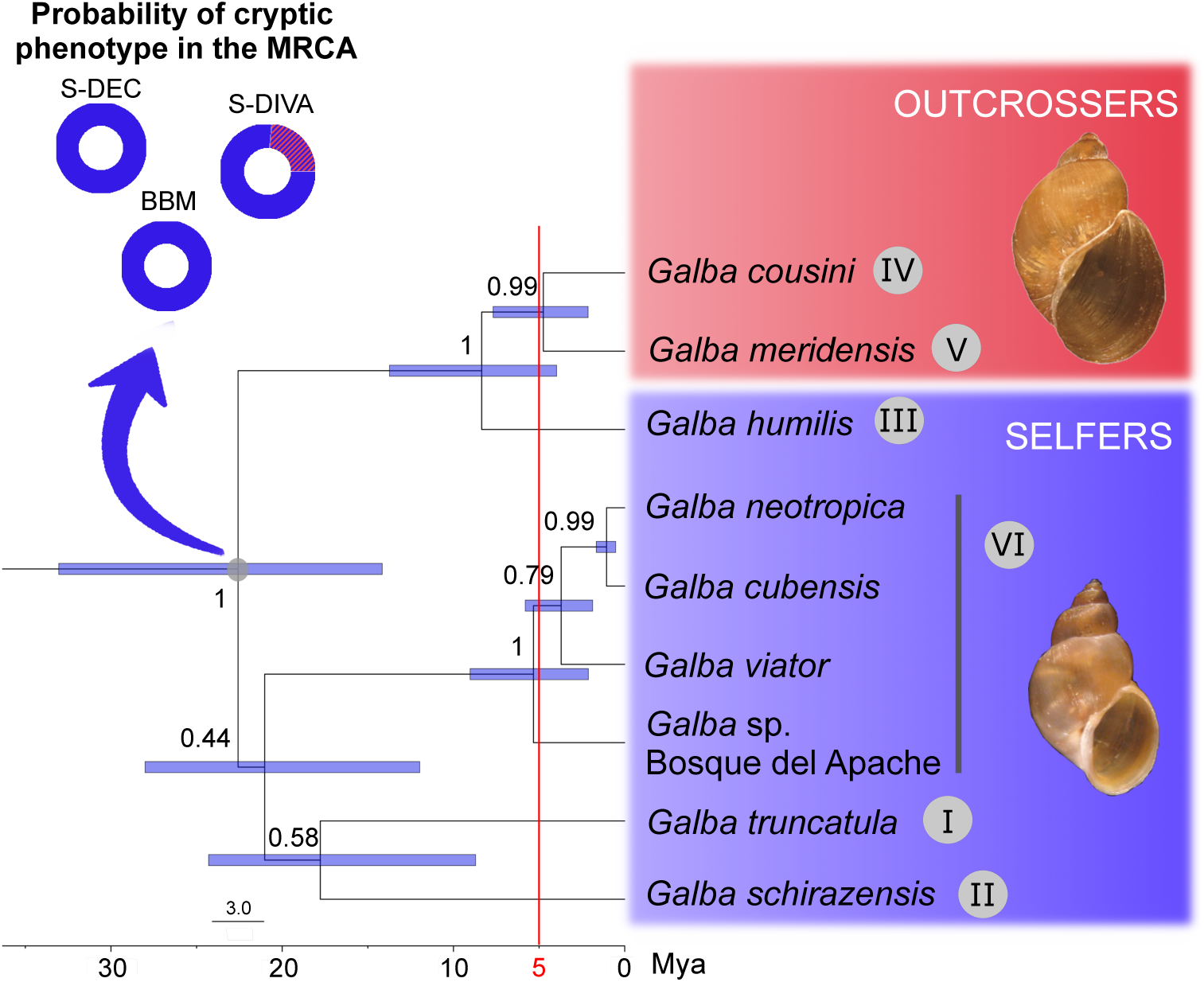
Most common topology of the species tree and phenotype of the most recent common ancestor of *Galba* inferred in StarBeast2. The probability of a cryptic phenotype in the most recent common ancestor of *Galba* species is 100% according to S -DEC and BBM and 78% according to S-DIVA. Clusters as appeared in text are shown in roman numbers. Node values indicate posterior probability and blue bars indicate 95% credibility intervals. The mating system is indicated on the right.

The estimated divergence time from the most recent common ancestor of the *Galba* group was 22.6 Mya [95% HPD interval: 14.6–33; Figs. 3–4]. Diversification within the *G. cousini*/*meridensis* species complex and the *G. cubensis*/*viator* species complex seems to have occurred 5 Mya, or less. The three phenotypic state reconstruction analyses (S-DEC, S-DIVA, and BBM) suggested that the most recent common ancestor of *Galba* species displayed the cryptic phenotype (Fig. 4). Thus, the phenotype of *G*. *cousini* and *G. meridensis* should be considered as derived from the cryptic phenotype.

## Discussion

### *Galba* comprises six clusters

Here we report the largest study published to date of *Galba* systematics and distribution, based on extensive sampling at a very large geographical scale and integration of phenotypic and molecular approaches across all DNA sequences available in GenBank for four genes (Dayrat 2005). The widespread occurrence of self-fertilization in these populations voids the biological species concept (Coyne and Orr 2004), and the absence of any reliable morphological distinction obviates the typological one, with the exception of *G. cousini*/*meridensis*. Thus, we are left with a phylogenetic approach, which suggests the existence of six clusters, perhaps corresponding to as many as eight species, or as few as five. These findings reinforce several previously-published works that have involved fewer genes and smaller sample sizes (Correa et al. 2010, 2011; Bargues et al. 2011b, 2011a; Standley et al. 2013; Lounnas et al. 2017).

We suggest that three of our clusters are best understood as one species each. The oldest taxonomic names for these are *Galba truncatula* (Müller 1774) for cluster I, *Galba schirazensis* (Küster 1862) for cluster II, and *Galba humilis* (Say 1822) for cluster III. The other three clusters (IV to VI) may in fact correspond to just two highly-diverse species or species complexes.

Cluster IV (*Galba cousini* Jousseaume 1887) and cluster V (*Galba meridensis* Bargues, Artigas, Khoubbane & Mas-Coma 2011) may be united into a single species complex, with *cousini* the oldest name. All these populations inhabit northern regions of South America, and together they exhibit a single phenotype not shared by other populations of *Galba*. They always clustered together in our phylogenetic reconstructions, with a divergence time of 4.7 Mya. However, *G. meridensis* has been sampled in but a single locality, and more extensive sampling is required to ascertain whether these are two species or a single species complex.

Cluster VI, including the nomina *Galba viator* (d’Orbigny 1835), *Galba cubensis* (Pfeiffer 1839), *Galba neotropica* (Bargues, Artigas, Mera y Sierra, Pointier & Mas-Coma, 2007), and a population from southern USA (Bosque del Apache) seem to comprise *Galba* species number five. The distance separating *G. cubensis* and *G*. *neotropica* is limited (1 Mya). Moreover, microsatellite markers defined in *G. cubensis* amplified effectively in individuals of *G*. *neotropica* (Lounnas et al. 2017), suggesting a very short genetic distance. We therefore suggest synonymy of these two species under the name *G. cubensis*. *Galba viator* and *G*. *cubensis* have overlapping distributions in Argentina, Chile, and Uruguay (Artigas et al. 2011; Sanabria et al. 2013; Standley et al. 2013; Medeiros et al. 2014), and additional sampling in this region would help resolve their genetic relationships. Here any distinction between the two sets of populations depends on both the genes and phylogenetic methodology employed. The population from Bosque del Apache (USA) was also included in cluster VI, depending on the gene analyzed. Its long mitochondrial branch may suggest rapid evolution, and here again more extensive sampling is required before reaching any conclusions. On the whole, a cautious position would be to suggest that cluster VI corresponds to a species complex or a species with wide diversity, as has been found in other freshwater snails from the clade Hygrophila (e.g., Ebbs et al., 2018; Mavárez et al., 2002; Pfenninger et al., 2006). We also note that if we ultimately recognize a single species; its name should be *viator*, and not *cubensis*, based on prior description.

The *Galba* species tree that we constructed based on a multispecies coalescent model returned three groups: one group uniting *G. truncatula* and *G. schirazensis*, another uniting *G*. *humilis* and *G*. *cousini/meridensis* and the last with *G*. *cubensis*/*viator*. This result partially agrees with some gene trees published in previous work (Correa et al. 2010, 2011; Bargues et al. 2011a). However, previous trees were based on single genes and smaller sample sizes. Our phylogenetic analysis also revealed some distinctive branches, including the mtDNA sequences from Ethiopia (Dayrat et al. 2011) and the samples from Bosque del Apache. These results may suggest undetected species or accelerated molecular evolution (Fourdrilis et al., 2016; Pinceel et al., 2005; Thomaz et al., 1996).

In North America, Burch (1982) recognized 22 species of small, mud-dwelling lymnaeids, which he grouped into the genus *Fossaria* with two subgenera, *Fossaria* (*s*.*s*.) with 11 species and *Bakerilymnaea* with 11 (see Table S5 for species names). Johnson et al. (2013) transferred these species to the genus *Galba*, but otherwise retained the Burch system. Included in the present analysis were topotypic samples of *obrussa* from Philadelphia and *parva* from Cincinnati, both of which we here show to be indistinguishable from topotypic *humilis*, collected at Owego, New York. Remigio (2002) contributed to Genbank a 16S sequence from a Canadian population he identified as *Fossaria obrussa*, also grouping with cluster III (*humilis*). We suggest that *obrussa*, *parva*, and the seven other uniquely North American specific nomina listed above ascribed by Burch (1982) to the subgenus *Fossaria* are junior synonyms of *G*. *humilis*, setting aside American populations of *G*. *truncatula* as distinct. In addition to his *obrussa* sequence, Remigio (2002) contributed a 16S sequence from Oklahoma to Genbank which he labelled “*Fossaria bulimoides*”. This sequence grouped with cluster VI (*cubensis*/*viator*) in our analysis. We suggest that all 12 specific nomina ascribed by Burch (1982) to the *Fossaria* subgenus *Bakerilymnaea* (Table S5) including *bulimoides* (Lea 1841), are junior synonyms of *G*. *cubensis*/*viator*.

### A set of cryptic species

Our study has confirmed the previous reports of Samadi et al. (2000) and Correa et al. (2011) that most species of *Galba* cannot be distinguished on the basis of shell morphology or internal anatomy. Trait variability within species seems to be greater than variance among species, likely attributable to phenotypic plasticity (Correa et al. 2011). Reproductive and growth traits in *G*. *truncatula* have been shown to vary according to habitat characteristics at small geographical scales (Chapuis et al. 2007), suggesting both that life-history traits are phenotypically plastic, and that to rely on such traits for specific identification is not advisable.

Our study also confirms that *G*. *cousini*/*meridensis* differs strikingly in adult size, shell shape, and anatomy from all other *Galba* species. Our molecular evidence suggests that *G. cousini/meridensis* evolved from a *Galba* ancestor demonstrating the cryptic phenotype. Interestingly, *G*. *cousini*/*meridensis* is the largest species within the genus *Galba*, occurs in a specialized habitat and displays the complex reproductive anatomy of an outcrossing species (see Jarne et al. 2010; Escobar et al. 2011).

Among the freshwater pulmonates, crypsis has previously been documented in *Ancylus* (Weiss et al. 2018) and *Radix* (Pfenninger et al. 2006). Our methods here were strictly qualitative, as was the case for *Ancylus* and *Radix*, because previous studies (Samadi et al. 2000; Correa et al. 2011) have shown that the dimensions of internal organs depend on physiological state and hence that species cannot be distinguished by means of such measurements. Future, more comprehensive approaches should include other more discrete anatomical traits, such as radular morphology.

Four hypotheses have been offered to explain the occurrence of cryptic species: recent divergence, parallelism, convergence, and stasis (Fig S1, Bickford et al. 2007; Fišer et al. 2018; Struck et al. 2018). The recent divergence hypothesis seems unlikely in this case. *Galba* has no closely related groups; its closest relatives are probably the stagnicoline lymnaeids of North America and Eurasia, which demonstrate a very distinctive morphology (Correa et al. 2010). Our analyses suggest that the several species of *Galba* are separated by more than 20 Myr (Burgarella et al. 2015). And indeed, the morphological divergence demonstrated by *G*. *cousini*/*meridensis* suggests that time is not been a significant constraint. The parallelism hypothesis also seems unlikely given that, based on our phylogenetic reconstruction, the cryptic morphology is ancestral for *Galba*, and the only other morphology that has evolved in the group, as demonstrated by *G*. *cousini*/*meridensis,* is derived. Nor does the topology of lymnaeid phylogeny fit the convergence hypothesis (Correa et al. 2010). So, by default, morphological stasis is left as the most likely hypothesis to explain crypsis in the genus *Galba*, as has been proposed in other gastropod groups (e.g., Gomez et al., 2004; Struck et al., 2018).

Strong stabilizing selection related to specialized environmental conditions is one possible explanation for the morphological stasis demonstrated by our several species of *Galba*. Populations have colonized habitats that are not exploited by other freshwater snails, especially the transiently inundated margins of water bodies exposed during the dry season(s) (see Table S1). In other words, *Galba* populations are more amphibious than other freshwater snails, which might mitigate both predation and interspecific competition (Burgarella et al. 2015). Adaptation to such habitats might impose strong stabilizing selection for a shell morphology able to resist desiccation and concomitant morphological stasis.

Among the many adaptations that have been correlated with life in transitory environments such as the exposed margins of water bodies is the evolution of self-fertilization (Escobar et al. 2011; Burgarella et al. 2015). The stasis in reproductive anatomy we have documented in populations of *Galba* may simply reflect a shared adaptation to self-fertilization in unpredictable habitat patches (Jarne et al. 2010). Once established, self-fertilization promotes the rapid erosion of genetic variation, as inbred populations become progressively unable to generate new genetic combinations through recombination (Noël et al. 2017). Thus, selfing could reinforce morphological stasis that might have evolved for other reasons.

The challenge of identifying cryptic *Galba* species is aggravated by their wide and poorly-known geographical distributions, recently scrambled by biological invasion. For example, *G*. *schirazensis* and *G*. *truncatula* have broadly expanded their distribution over recent decades (Brown 1994; Bargues et al. 2001, 2011a; Vinarski and Kantor 2016; Lounnas et al. 2018). We have documented up to three *Galba* species occurring in some South American sites (Table S1). Their identification is not possible without molecular tools.

The specific identity of *Galba* populations is important because they are involved in the transmission of fasciolosis caused by the liver fluke *F*. *hepatica*. Some studies have shown that lymnaeid species demonstrate different patterns of susceptibility, host-parasite compatibility and immunological resistance to *F*. *hepatica* (Gutiérrez et al. 2003; Vázquez et al. 2014; Dreyfuss et al. 2015). Although all species can be infected under laboratory conditions (Vázquez et al. 2018), field transmission depends on ecological and sociological conditions. Cattle or wildlife do not occupy the same grazing habitats as infecting snails in many parts of the world (Sabourin et al. 2018). Ecological studies should be performed to evaluate whether the several cryptic *Galba* species differ with regard to habitat preference, since our current knowledge is essentially limited to *G*. *truncatula* (Chapuis et al. 2007).

### Conclusions and future directions

Lymnaeid populations of the genus *Galba* are of interest for addressing a variety of questions, including wide-scale biogeography, biological invasions, evolution of mating systems, and host-parasite interactions. Our work is a first attempt to clarify the phylogeny, systematics, and biogeographical distribution of this interesting group in the New World. We have constructed a variety of gene trees using classical approaches, as well as a species tree based on a multispecies coalescent model that reconciles gene trees and provides a much better estimation accuracy for species tree topology than, for instance, concatenation (Heled and Drummond 2010). The inferred phylogenetic relationships among species varied, depending on the genes analyzed and techniques employed. Such a discordance is not unusual and has been reported in many different studies (e.g., Kutschera et al., 2014; Stewart et al., 2014; Suh et al., 2015), including in mollusks (Krug et al. 2013; Sales et al. 2013). Incomplete lineage sorting or introgressive hybridization of specific genes may indeed lead to such a result (Felsenstein 2004). Future studies could investigate which evolutionary processes (gene duplication, horizontal gene transfer, incomplete lineage sorting, hybridization) gave rise to the incongruence we have observed in gene and species trees.

The integration of fossil data would also likely provide more accurate insights into macroevolutionary dynamics of *Galba* than molecular phylogenies alone. The fossil record has proven very helpful in determining the phylogeny of groups containing hard body parts like plants, mammals, and mollusks (Magallón and Sanderson 2005; Agnarsson et al. 2011; Bolotov et al. 2016). But unlike marine gastropods that have a diverse fossil record with many helpful shell characters (sculptures, ornamentation, coloration, protoconch; Bouchet and Strong 2010), fossil freshwater pulmonates are scarce and lack useful taxonomic characters. The fossils that have been attributed to *Galba* are not strikingly distinct from those of other species belonging to other genera of Lymnaeids. Baker (1911) reported *Galba*-like fossils in the Cretaceous of North America, but these shells might also belong to other genera of Lymnaeids, or even an ancestor of current Lymnaeids. When the generalized shell morphology is added to the absence of anatomical data, the application of fossil data to *Galba* phylogenetic reconstruction becomes excessively problematic.

Although our study was conducted at an extremely large geographic scale, especially in America, *Galba* populations occur on almost all continents. Much more extensive sampling and molecular analysis will be required to get a worldwide picture of the phylogeny and distribution of the genus. Of special interest is North America, where we have confirmed the occurrence of *G*. *humilis*, *G*. *cubensis*/*viator*, and *G*. *schirazensis*. We did not, however, confirm *G*. *truncatula*, despite its otherwise worldwide distribution (Bargues et al. 2001; Correa et al. 2011; Vinarski et al. 2011; Novobilský et al. 2014). The recent arrival of *G*. *schirazensis* in Europe and the Middle East (Bargues et al. 2011a; Lounnas et al. 2018) and the recent description of *Galba robusta* from Yemen, based on shell and penial morphology alone (Vinarski 2018) should be confirmed with molecular approaches.

Broader, worldwide sampling will be required to (i) confirm that the European, African and Asian populations referred to as *G*. *truncatula* actually belong to this species, (ii) clarify the biogeographic origin of *G. truncatula*, (iii) evaluate whether the *G. cubensis*/*viator* and *G. cousini*/*meridensis* groups constitute species complexes or single species with wide diversity, (iv) understand how the invasive species (especially *G*. *truncatula* and *G*. *schirazensis*) are spreading, and (v) confirm the loss of genetic variation in those populations as reported by other authors (Meunier et al. 2001; Lounnas et al. 2018).

It might also be useful to construct maps of *Galba* absence, possibly associated with detection probabilities. For example, *Galba* has not been found in Northern Brazil (Paraense 1982), possibly because the acidic waters of the Amazon and Orinoco rivers have blocked colonization (Paraense 1982, 1983). *Galba* populations are also absent from the most southern regions of Argentina (authors’ unpublished data). Negative records in species distribution can be important when constructing accurate distribution maps and inferring species-environment associations (Brotons et al. 2004). Models based only on presence data are less predictive (Václavík and Meentemeyer 2009). Absence data have been used to good effect modelling the spatial distribution of the medically important freshwater snail species *Bulinus globosus*, *Biomphalaria pfeifferi*, and *Radix natalensis* in Zimbabwe and predicting their distribution in a future climate (Pedersen et al. 2014).

More detailed ecological studies, based on long-term surveys of sites and analyses of life-history traits under laboratory conditions (e.g., Chapuis et al., 2007) would facilitate our understanding of species interactions and the transmission of *F*. *hepatica* (Sabourin et al. 2018). For example, *G*. *schirazensis* seems to be spreading very efficiently and genetic studies suggest that this is mainly due to one genetic clone (Lounnas et al. 2018). Studying the competitive and transmissive abilities of this clone, compared to both other *G*. *schirazensis* and other *Galba* species would be worthwhile.

## Supporting information

Supp. Mat. Figure S1

Supp. Mat. Figure S2

Supp. Mat. Figure S3

Supp. Mat. Figure S4

Supp. Mat. Figure S5

Supp. Mat. Figure S6

Supp. Mat. Figure S7

Supp. Mat. Figure S8

Supp. Mat. Figure S9

Supp. Mat. Figure S10

Supp. Mat. Figure S11

Supp. Mat. Figure S12

Supp. Mat. Figure S13

Supp. Mat. Tables S1-S5

Supp. Mat. References

## Data and code accessibility

Xml files for phylogenetic analyses are available from the Zenodo repository (https://zenodo.org/record/3473937#.XZiPcC0ryTd).

## Acknowledgments

We would like to express our gratitude to Nicolás Bonel for useful comments on earlier drafts of the manuscript and Harry G. Lee for advice and assistance on the taxonomy. We thank Jimena Guerrero, Björn Stelbrink and Thomas Wilke for suggestions on phylogenetic analyses and for Graham R. Jones, Patricio Maturana Russel and Remco R. Bouckaert for assistance in running STACEY and Multi-threaded nested sampling. We thank the reviewers Pável Matos-Maraví and Christelle Fraïsse and the recommender from PCI in Evolutionary Biology Fabien Condamine for their thoughtful comments and suggestions. Fellowships granted by Erasmus Mundus PRECIOSA and Méditerranée Infection supported research stays of PA at the Institute de Recherche pour le Développement, MIVEGEC (Montpellier, France). AV was supported by a pour le Développement grant from IRD (BEST) and ML by a doctoral fellowship from University of Montpellier and a post-doctoral grant from Labex CeMeb. This study was financially supported by IRD, CNRS, ECOS-SUD (A16B02) and Malacological Society of London. Version 3 of this preprint has been peer-reviewed and recommended by Peer Community In Evolutionary Biology (https://doi.org/10.24072/pci.evolbiol.100089).

## Supplementary Material

### Table legends

**Table S1.** Sampled sites from America in which *Galba* species were found. Individuals were submitted to the three-step procedures for species identification (see text and Fig. 1). For each site, we provide the country, site name, geographic coordinates, sampling date, and number of sampled individuals. Note that only a fraction of sampled individuals was sequenced. For each step (and species), we indicate the number of individuals considered. NA: not available. * indicate sites that have been resampled at different dates. Accession names in GenBank (ITS2 and COI) are indicated into parentheses. Note that in some cases a single sequence was obtained. The last column show the 16S and ITS1 sequences that have been incorporated to the study in order to test the species hypothesis with the multispecies coalescent models.

**Table S2.** Sites retrieved from literature and GenBank where *Galba* species were molecularly identified in America. Both the *Galba* and *Lymnaea* names have been used in the literature at genus level for the species considered in our study—we used *Galba* here for this monophyletic group of small lymnaeids. For each site, we report the country, site, geographical coordinates available sequences of mitochondrial (COI and 16S) and nuclear (ITS1 and ITS2) genes, species identification by specific microsatellites, bibliographic reference, and the species name used in the reference. Coordinates from Owego, New York were obtained from GoogleEarth and those from Correa et al. (2010) from Correa et al. (2011). Some coordinates were corrected in order to match the specific site: Rio Negro (Argentina) from Correa et al. (2010), Frias (Argentina) from Correa et al. (2011) and Lounnas et al. (2017a), Estanque Lagunillas (Venezuela) from Bargues et al. (2011c), Baños del Inca (Peru) from Bargues et al. (2012), Paysandú (Uruguay) from Lounnas et al. (2017a) and Geffrier (Guadeloupe) (provided by the authors). The KT461809 sequence was erroneously tagged as an ITS2 sequence, but is, in fact, a COI sequence. Sequences of the individuals molecularly identified by (Medeiros et al. 2014) are missing in the original publication and were not uploaded to GenBank. ND, no data available.

**Table S3.** Sites retrieved from literature and GenBank where *Galba* species were molecularly identified in Europe, Asia, and Africa. Coordinates that were not given in the original articles or in GenBank were best-guess estimated. The information reported for each site is as in Table S2. ND, no data available.

**Table S4.** Nested sampling results for the eleven species-delimitation models shown in Figure S13. The model with the higher Marginal Likelihood estimate is the top-ranked model. All Bayes factor (BF) calculations are made against the current taxonomy model (scenario D). Therefore, negative BF values indicate support for the current taxonomy model, and positive BF values indicate support for the alternative model.

**Table S5.** Species of small, mud-dwelling lymnaeids recognized by Burch (1982) in North America. The author grouped the 22 species into the genus *Fossaria* with two subgenera, *Fossaria* (*s*.*s*.) and *Bakerilymnaea*.

### Figure captions

**Figure S1.** Four evolutionary processes that can lead to cryptic species illustrated by their phylogenetic histories. (A) Recent divergence: cryptic species are very closely related and only recently diverged from each other and, thus, the rate of morphological disparity between them (A1 and A2) is not substantially different from that for non-cryptic species. (B) Parallelism: the cryptic species are not very closely related to each other and the rate of morphological disparity for non-cryptic species is much greater than that for cryptic species. (C) Convergence: the cryptic species are also not closely related to each other. Initially, morphological disparity for cryptic species can change in a manner similar to that for the non-cryptic species pair. However, at some point, morphological disparity decreases for the cryptic species, while continuing to increase between non-cryptic taxa. (D) Stasis: the cryptic species are closely related to each other or are part of a species complex and diverged a long time ago. In comparison with non-cryptic species, the rate of morphological change is substantially reduced. (Figure and legend extracted and modified from Struck et al. 2018).

**Figure S2.** First two planes of a principal component analysis on reproductive traits in Lymnaeid species. These planes represent 80.36% of the total variance. The analysis was conducted using the following measurements: prostate height, prostate width, penis length, preputium length, anterior and posterior widths of the penis sheath, and shell length and width. All variables were ln-transformed. This analysis included most of the species considered in our work, as well as *Pseudosuccinea columella* (one of the closest relative of *Galba* spp.), sampled in Argentina, Colombia, Cuba, France, Guadeloupe, Peru and Venezuela. Each point represents an individual (N = 81). (Figure and legend extracted and modified from Correa et al. 2011).

**Figure S3.** Geographic distribution of *Galba cubensis*, *Galba schirazensis* and *Galba truncatula* in the European, Asian and African samples retrieved from GenBank. Coordinates are given in Table S3.

**Figure S4.** Shells and reproductive and urinary systems of the six *Galba* species studied. *Galba cousini*/*meridensis* is the only species that can be morphologically identified.

**Figure S5.** Phylogenetic tree of *Galba* species based on Bayesian inference in Beast2 of the COI gene. All sequences from the current study, as well as the ones retrieved from GenBank, are included in this tree. Sequence coloration represents species. Arrows indicate sequences belonging to a type locality (see Table 1 for details). Sequence data are given in Tables S1, S2, and S3.

**Figure S6.** Phylogenetic tree of *Galba* species based on Bayesian inference in Beast2 of the 16S gene. All sequences were retrieved from GenBank except for sequences from individuals from Bosque del Apache (USA). Sequence coloration represents species. Arrows indicate sequences belonging to a type locality (see Table 1 for details). Sequence data are given in Tables S1, S2, and S3.

**Figure S7.** Phylogenetic tree of *Galba* species based on Bayesian inference in Beast2 of the ITS1 gene. All sequences were retrieved from GenBank except for sequences from individuals from Bosque del Apache (USA). Sequence coloration represents species. Arrows indicate sequences belonging to a type locality (see Table 1 for details). Sequence data is given in Tables S1, S2, and S3.

**Figure S8.** Phylogenetic tree of *Galba* species based on Bayesian inference in Beast2 of the ITS2 gene. All sequences for the current study, as well as the ones retrieved from GenBank, are included in this tree. Sequence coloration represents species. Arrows indicate sequences belonging to a type locality (see Table 1 for details). Sequence data are given in Tables S1, S2, and S3.

**Figure S9.** Haplotype network of *Galba* species based on 16S gene. Circle sizes are proportional to haplotype frequencies and colors represent species. The number of mutations separating circles are indicated by dashes. The six clusters detected in the phylogenetic analysis are represented as grey shapes.

**Figure S10.** Haplotype network of *Galba* species based on COI gene. Circle sizes are proportional to haplotype frequencies and colors represent species. The number of mutations separating circles are indicated by dashes. The six clusters detected in the phylogenetic analysis are represented as grey shapes.

**Figure S11.** Haplotype network of *Galba* species based on ITS1 gene. Circle sizes are proportional to haplotype frequencies and colors represent species. The number of mutations separating circles are indicated by dashes. The six clusters detected in the phylogenetic analysis are represented as grey shapes.

**Figure S12.** Haplotype network of *Galba* species based on ITS2 gene. Circle sizes are proportional to haplotype frequencies and colors represent species. The number of mutations separating circles are indicated by dashes. The six clusters detected in the phylogenetic analysis are represented as grey shapes.

**Figure S13.** Scenarios for species assignments used to run the multispecies tree models using Multi-Threaded Nested Sampling in StarBeast2. The scenario K is an unreal scenario that separates the populations of *G. viator* from Argentina and Chile to test whether splitter models showed higher support than lumper models regardless its biological sense.

