## Supplementary figures and images for "Systematics and geographical distribution of *Galba* species, a group of cryptic and worldwide freshwater snails"

### Supp. Mat. Figure S1

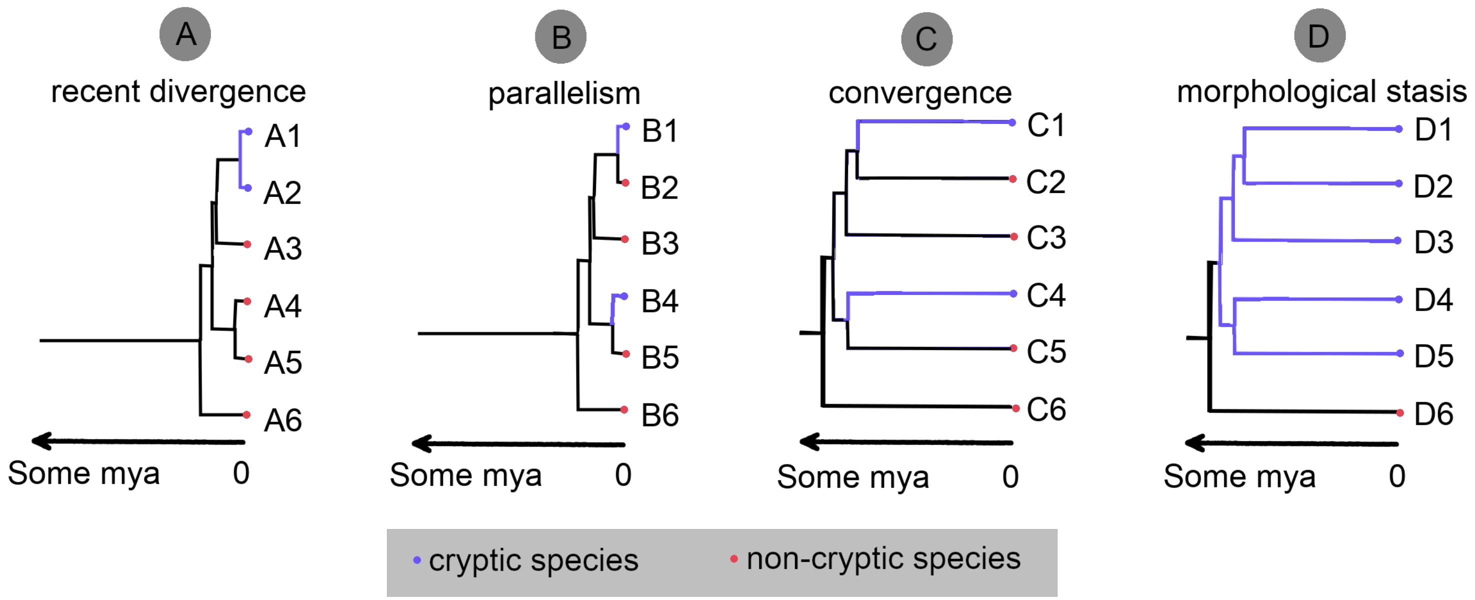

### Supp. Mat. Figure S2

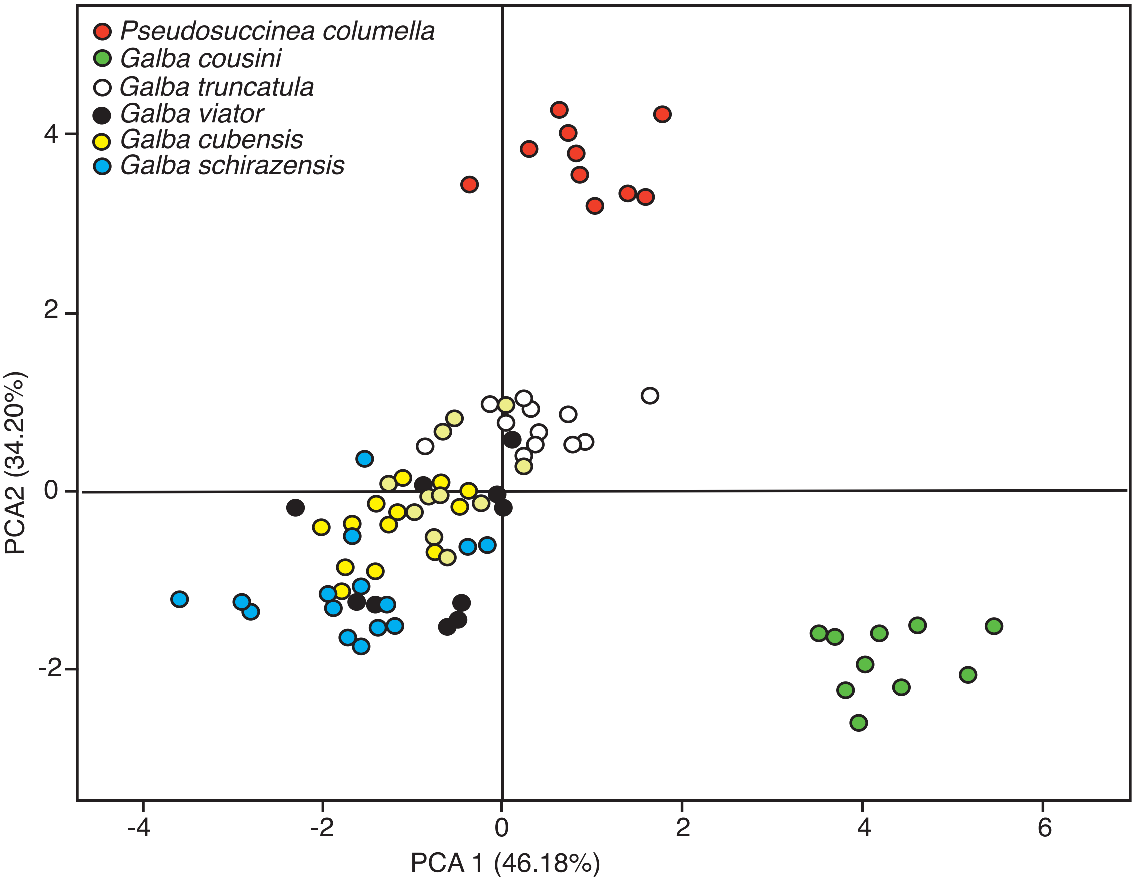

### Supp. Mat. Figure S3

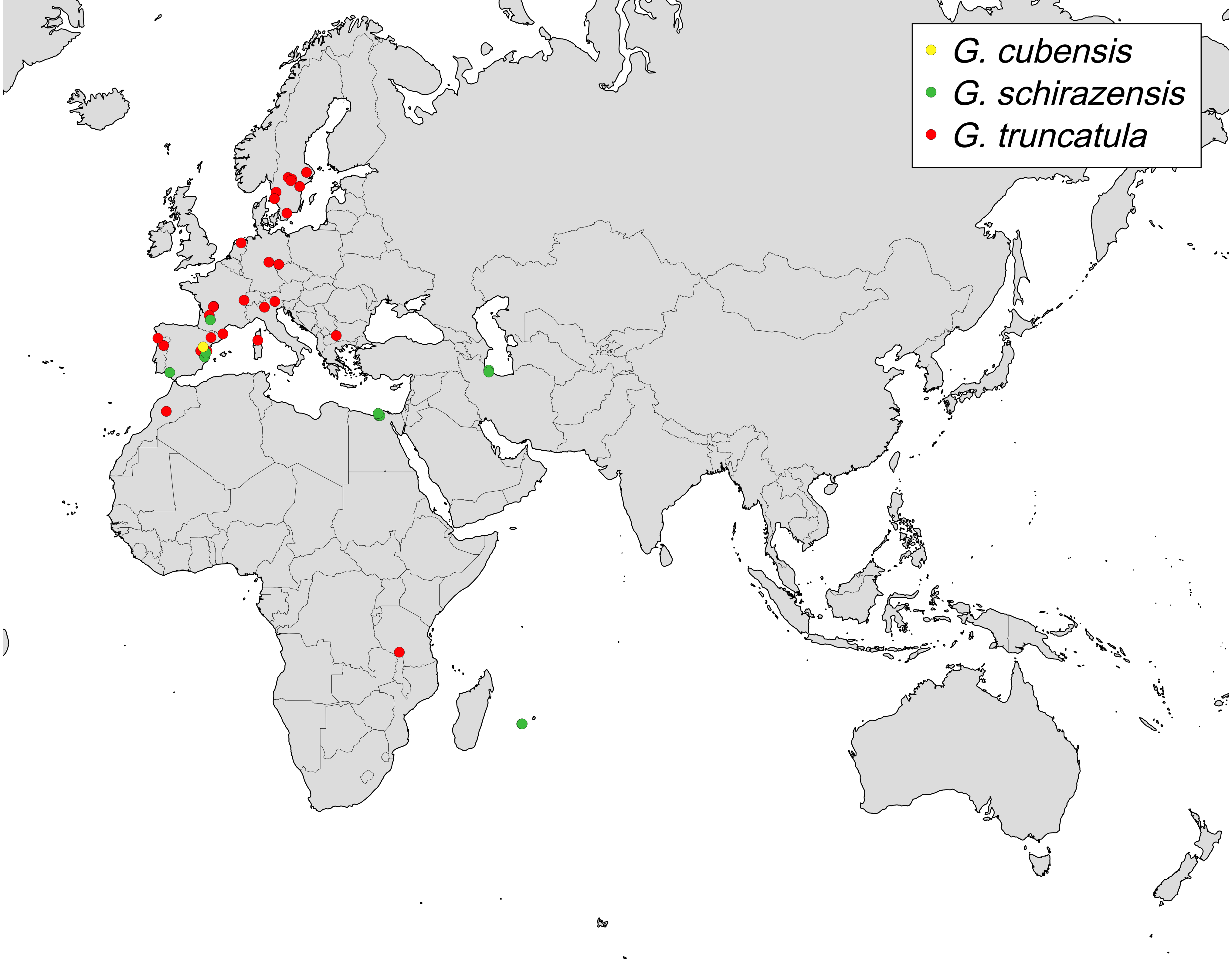

### Supp. Mat. Figure S4

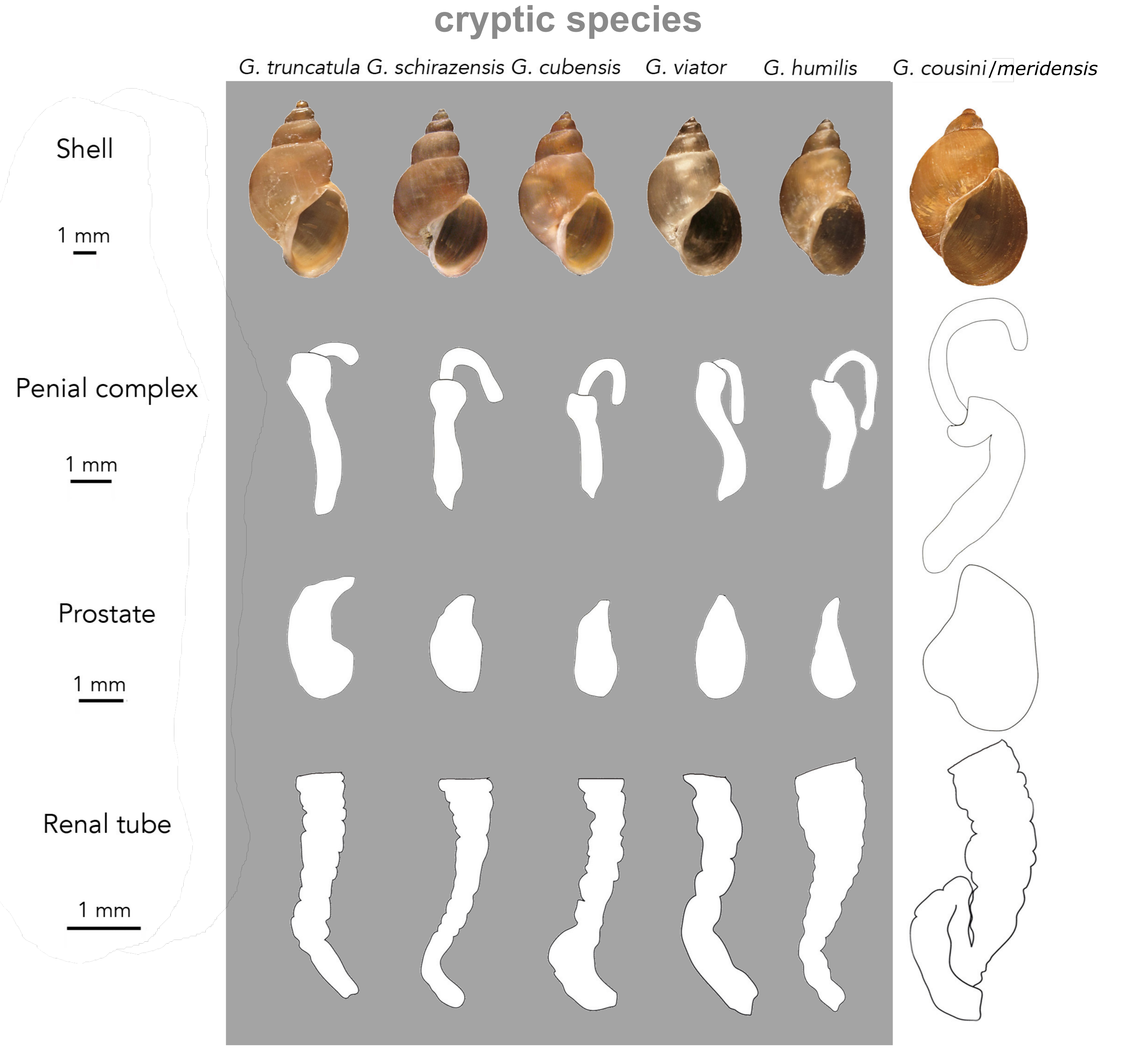

### Supp. Mat. Figure S5

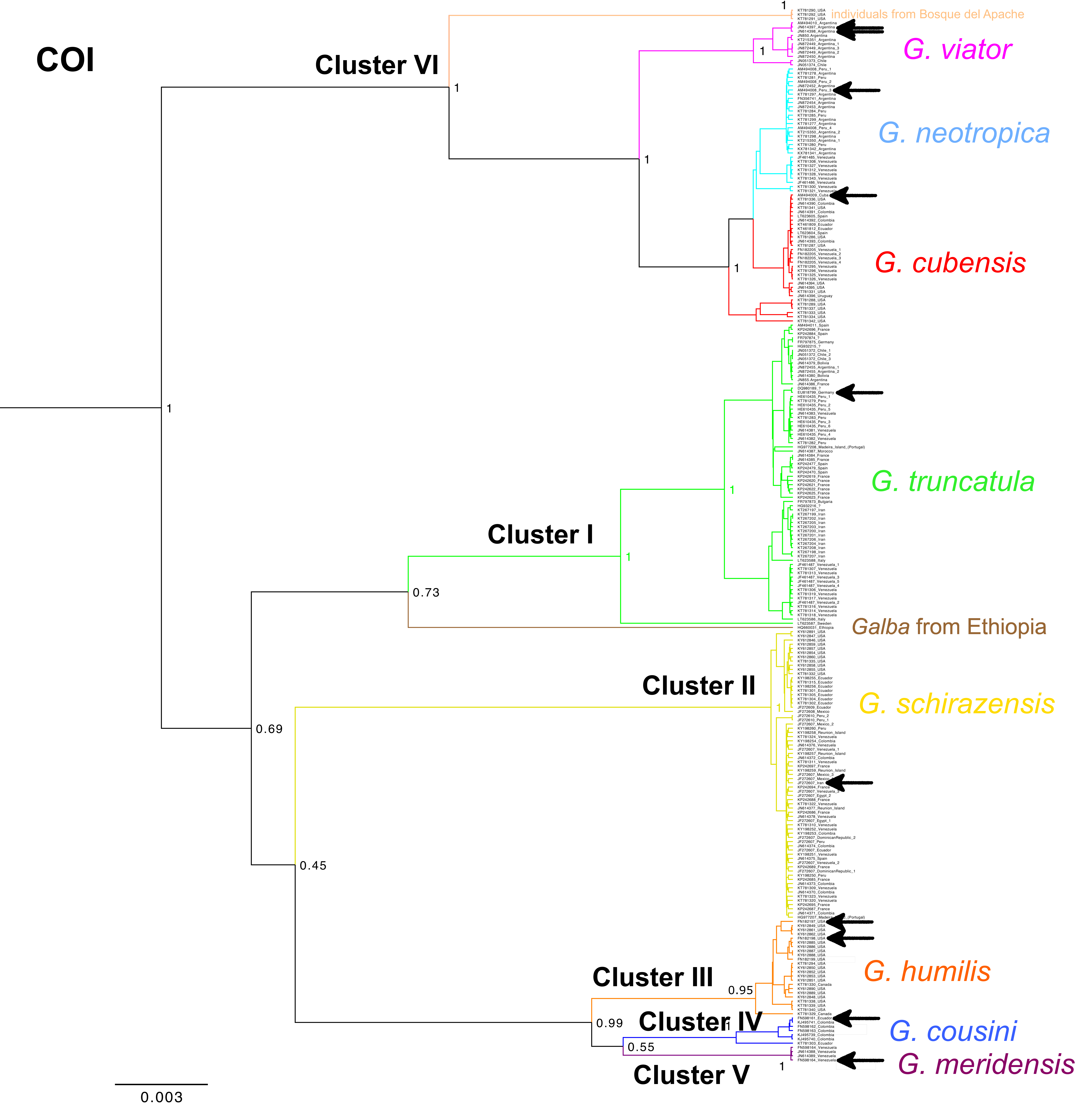

### Supp. Mat. Figure S6

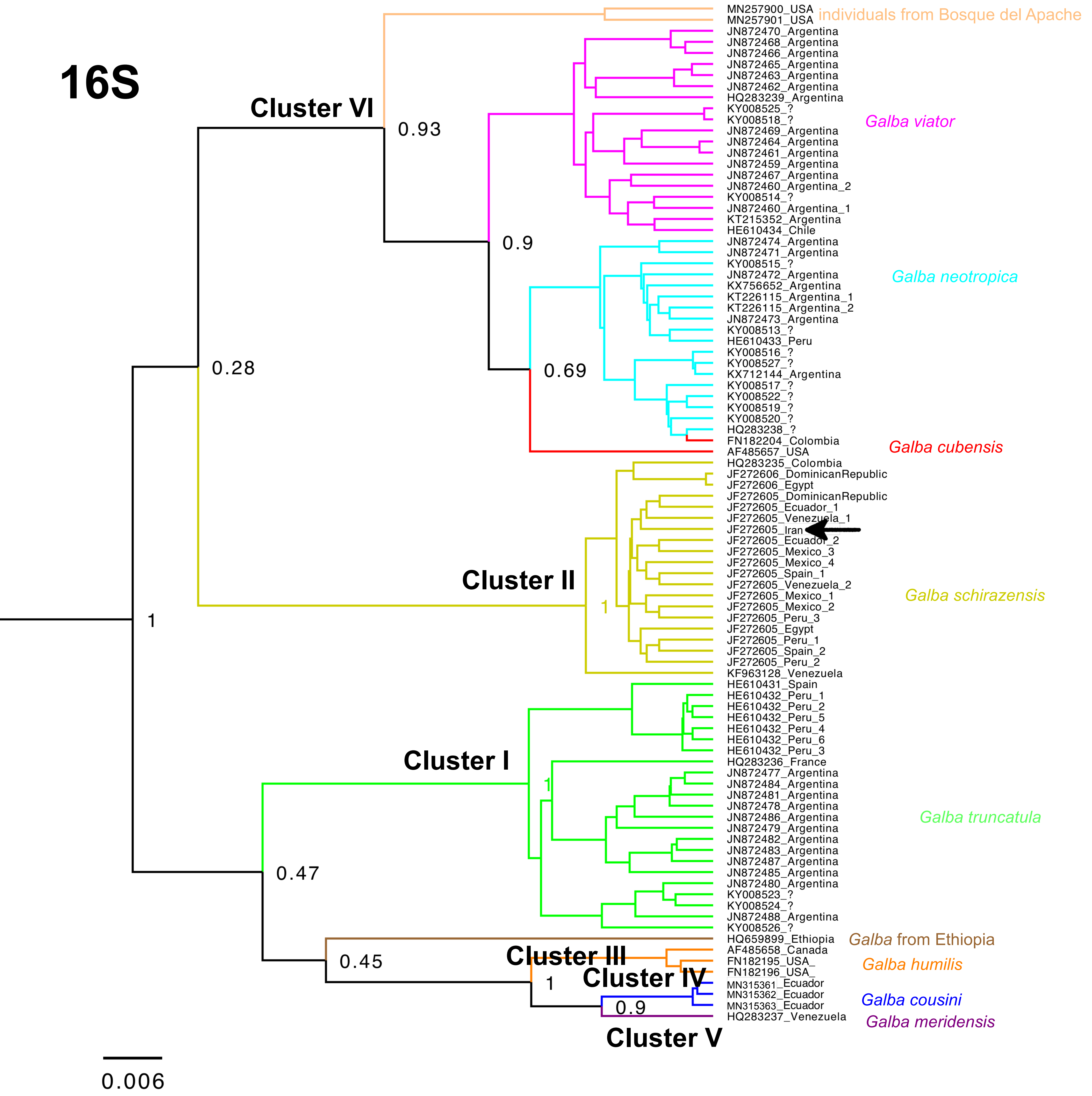

### Supp. Mat. Figure S7

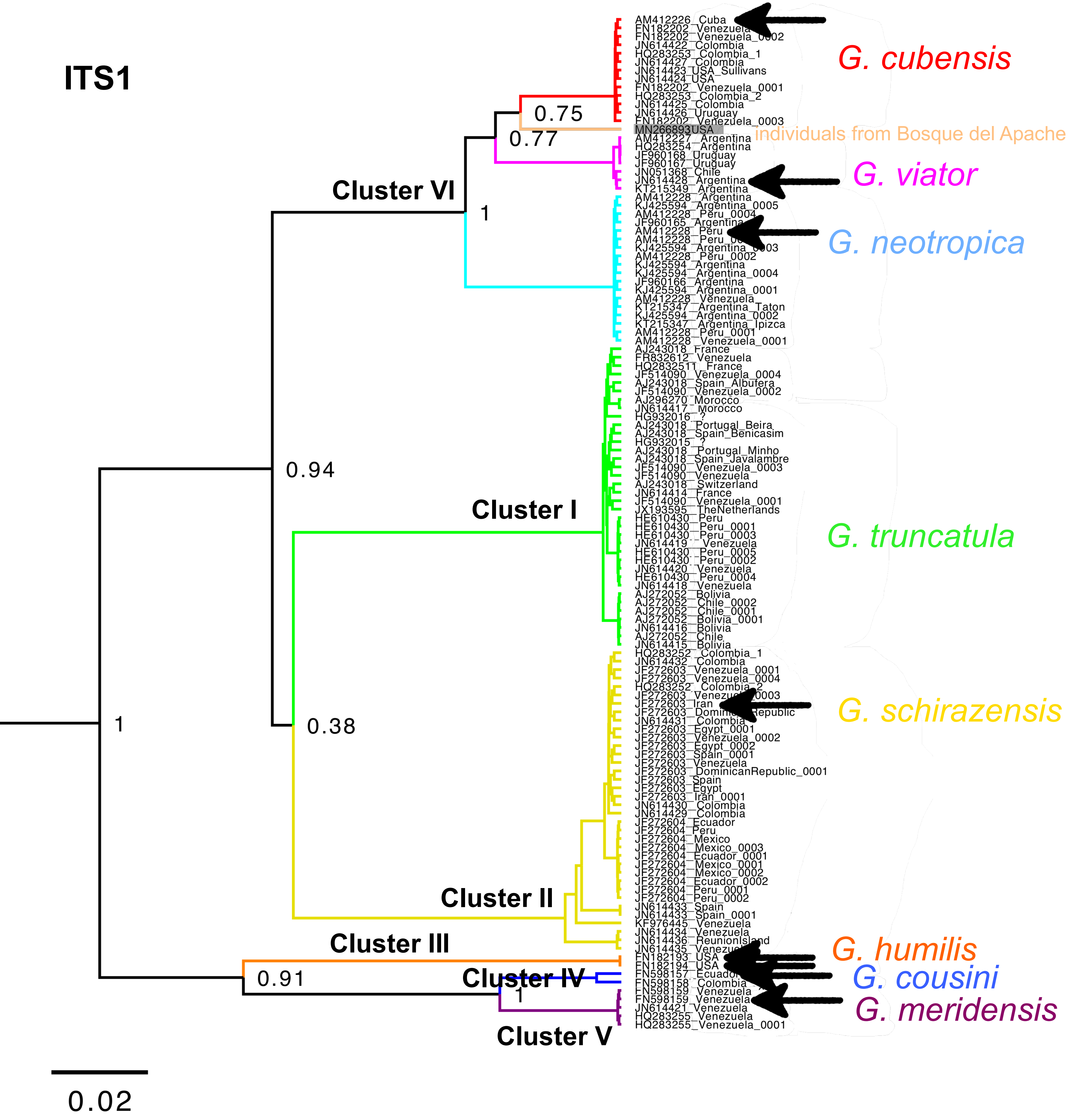

### Supp. Mat. Figure S8

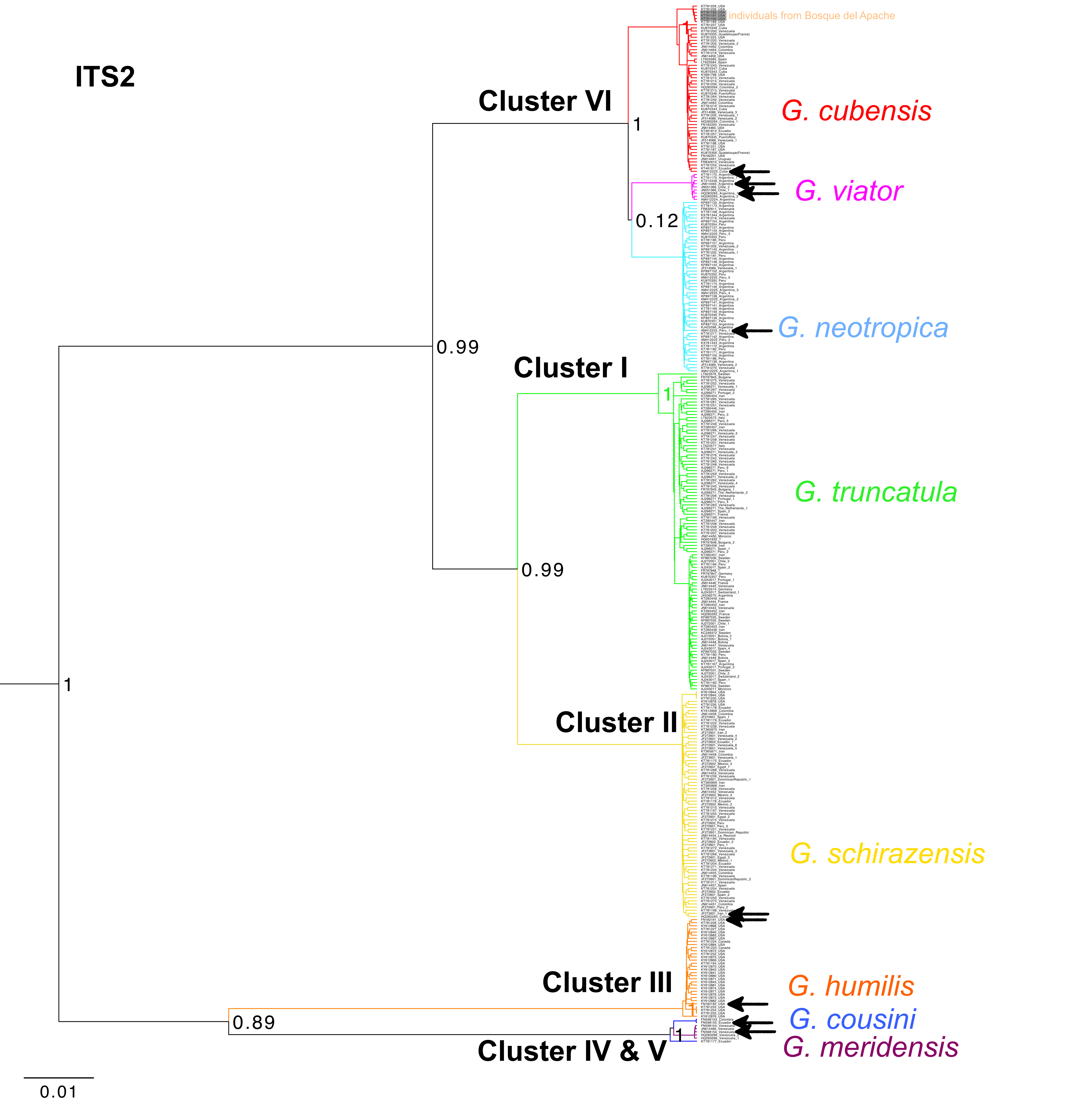

### Supp. Mat. Figure S9

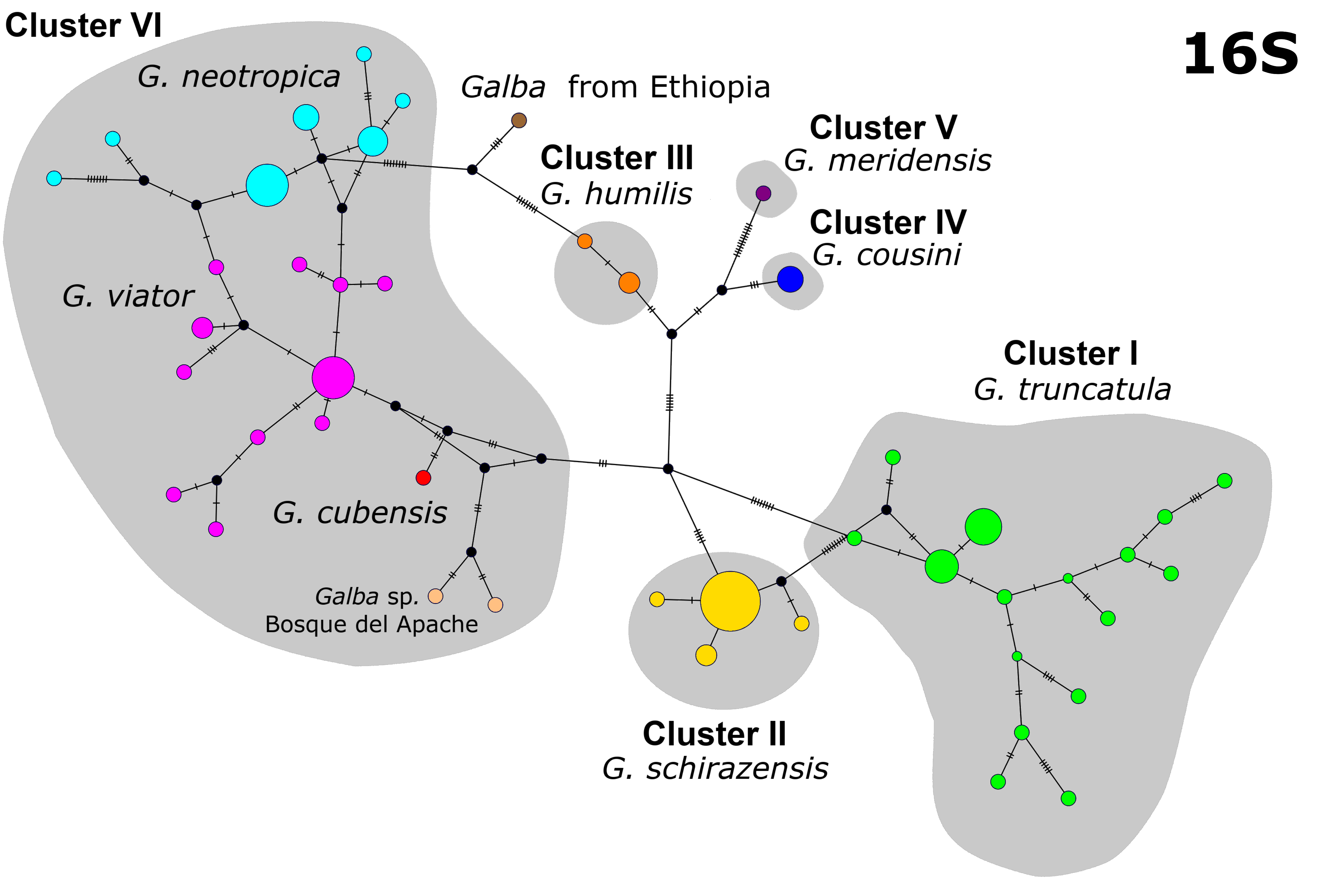

### Supp. Mat. Figure S10

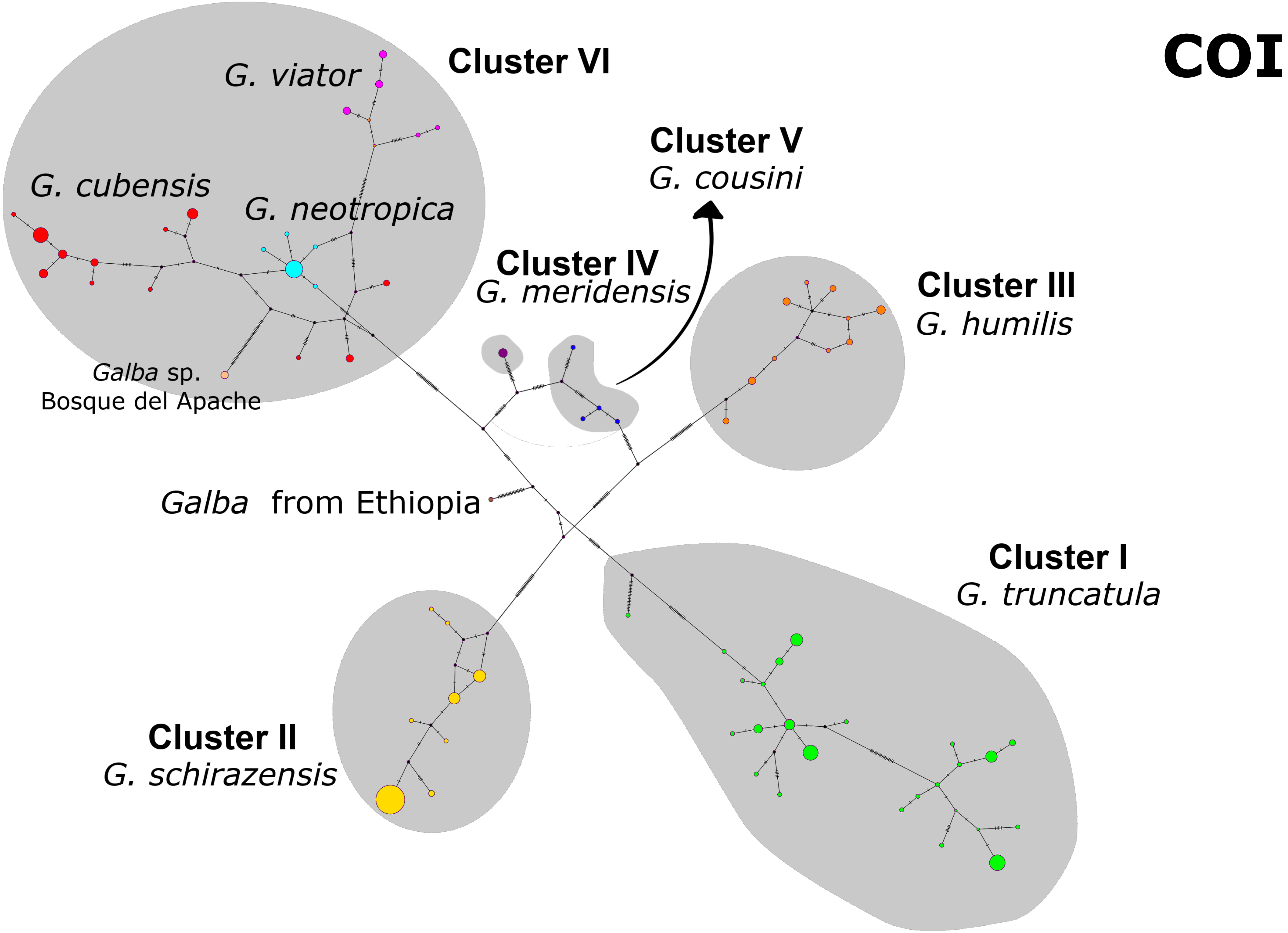

### Supp. Mat. Figure S11

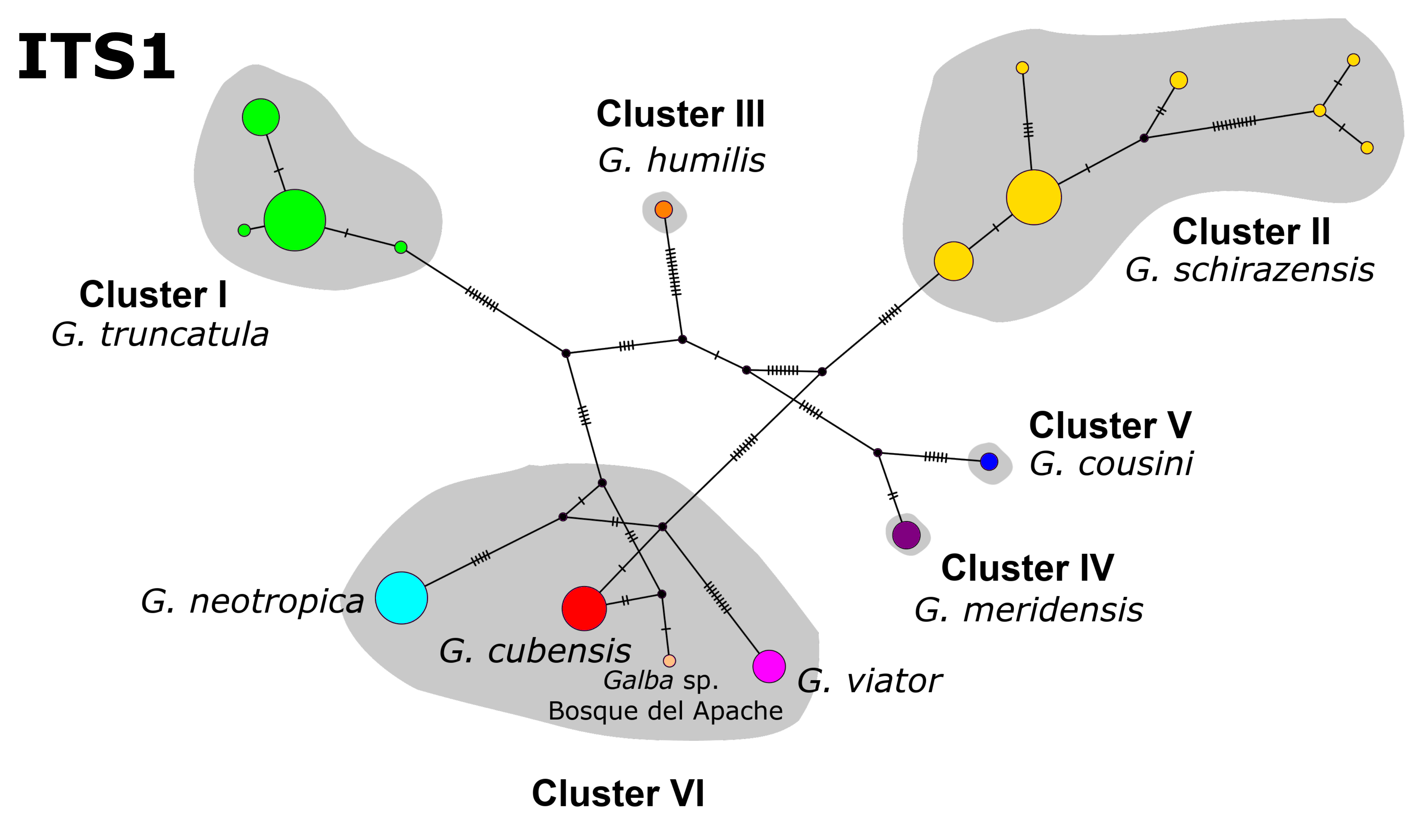

### Supp. Mat. Figure S12

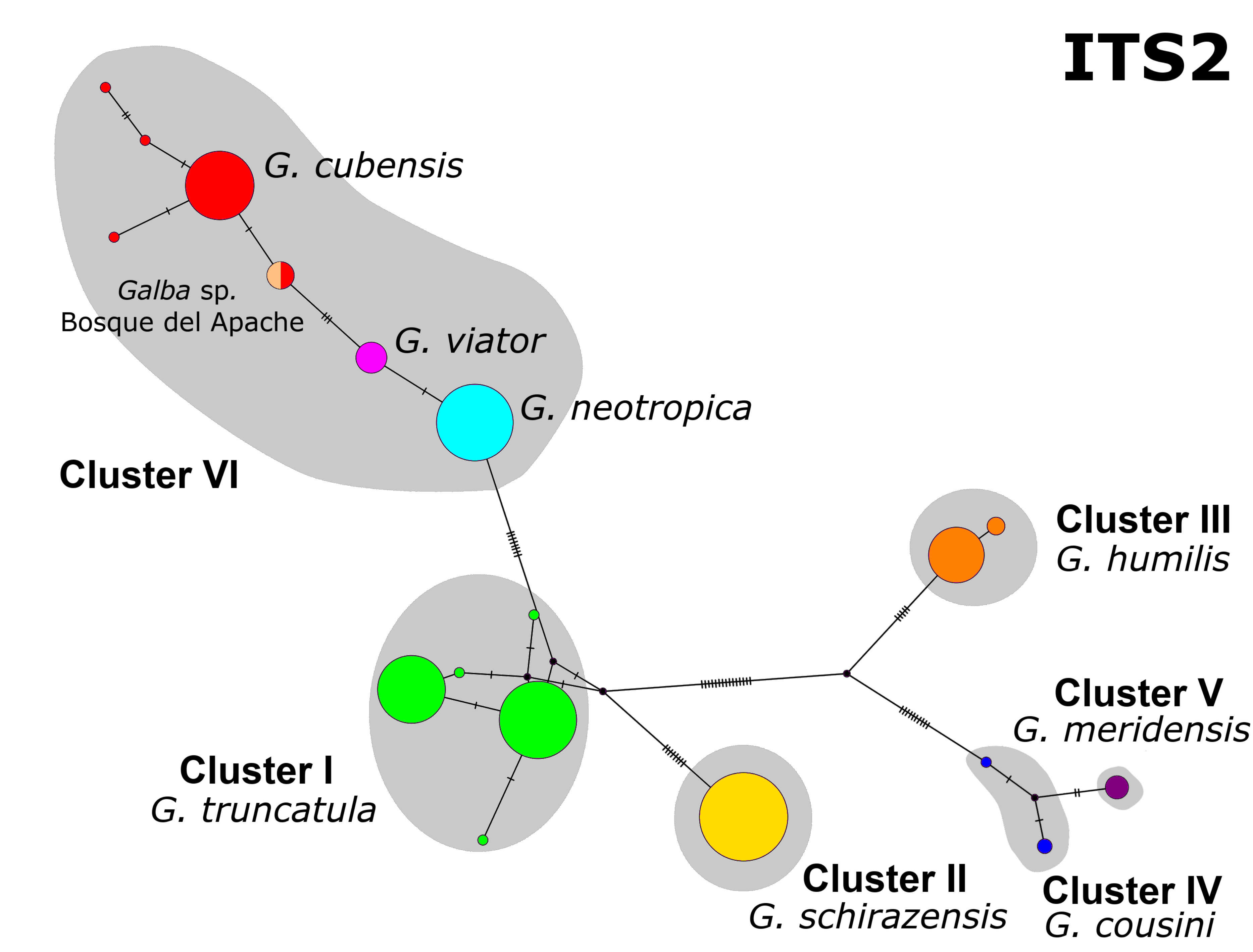

### Supp. Mat. Figure S13

#### Scenarios tested with Nested Sampling analysis

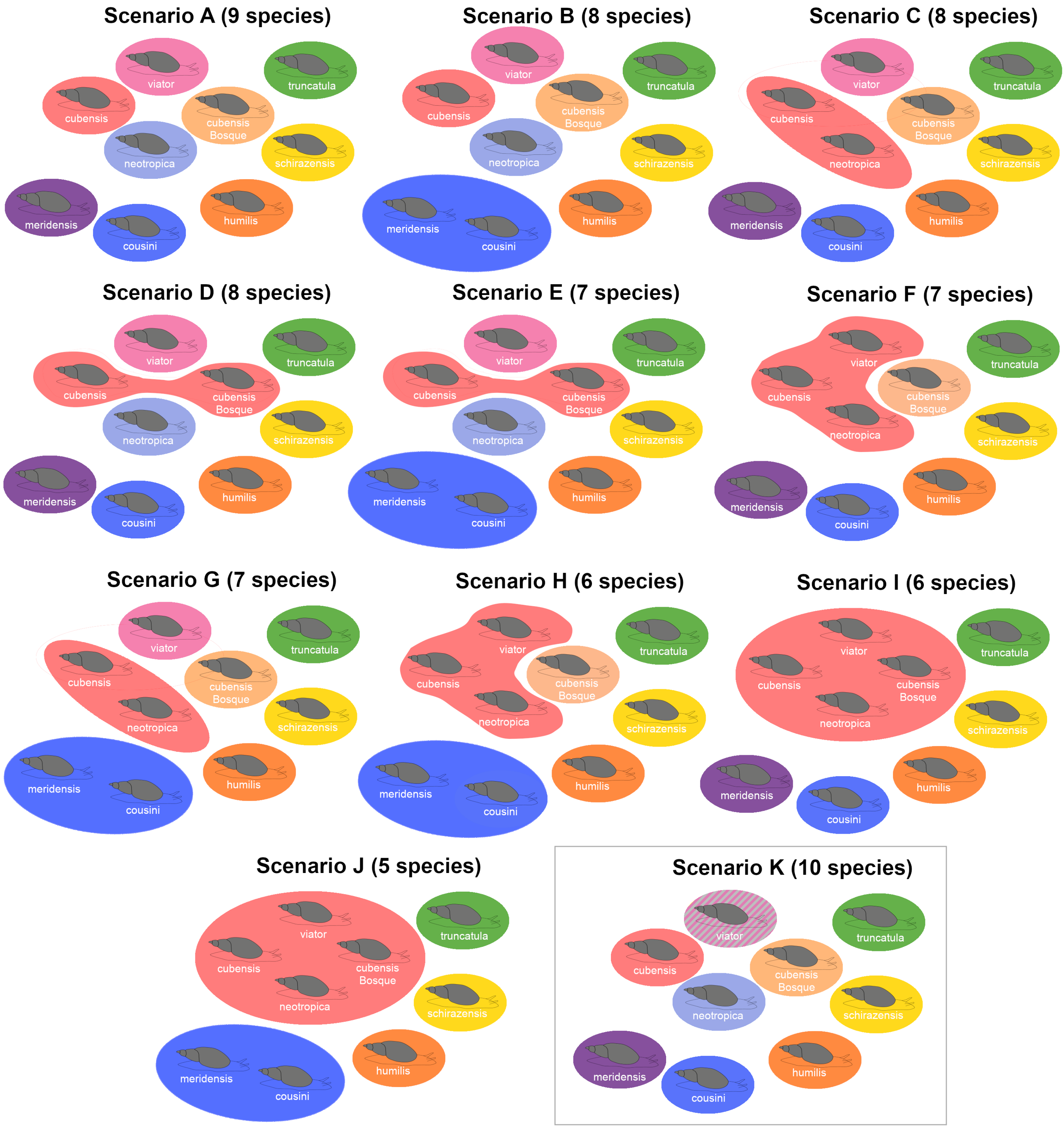
